# Promises and limitations of local ancestry inference in imputed ancient genomes

**DOI:** 10.64898/2026.05.19.725905

**Authors:** Katia Bougiouri, Evan K. Irving-Pease, Laurent A.F. Frantz, Fernando Racimo, Martin Petr

## Abstract

Recent advances in genome imputation have enabled the application of state-of-the-art statistical methods—originally developed for present-day genomes—to ancient genomes. One class of such methods, known as local ancestry inference (LAI), can model an individual’s genome as a mosaic of tracts assigned to different putative ancestral sources, revealing patterns of genetic ancestry across the genome. However, most LAI methods have been designed to study recent admixture events in human history, and they generally assume large panels of present-day genomes. Despite the recent availability of high-quality imputed ancient genomes, it remains unknown to what degree LAI inference is reliable for such datasets. Ancient DNA is often characterized by heterogeneous geographic and temporal sampling, varying degrees of divergence between ancient source proxies and admixing populations, and complex demographic histories. Here, we performed an extensive set of population genetic simulations to evaluate the accuracy of four popular LAI methods–RFMix, FLARE, MOSAIC and simpLAI–under different demographic scenarios, various temporal sampling schemes, sample sizes, and admixture dates. We quantify the accuracy of these methods as a function of different parameters in practically relevant scenarios, and provide general guidelines for future studies utilizing LAI in ancient DNA research.

## Introduction

Throughout evolutionary history, the process of recombination causes a rearrangement of genetic information, producing a complex mosaic of genealogical histories carried by each genome. The resulting patterns of genetic variation can be used to reconstruct different aspects of this history, including past episodes of admixture between ancestral groups. A popular class of methods consists of assigning or ‘painting’ haplotypes as originating from one of multiple putative source ancestries; a process also known as local ancestry inference (LAI). These methods commonly rely on providing reference populations as proxies for the true ancestral sources, and have been primarily developed to reconstruct ancestral histories of recently admixed genomes (Gravel 2012). LAI has not only served to track past migrations and time admixture events (Johnson et al. 2011; Baharian et al. 2016; Medina et al. 2018), but has also contributed to studies focusing on association testing while controlling for population structure (Diao and Chen 2012; Xu and Guan 2014; Zhong et al. 2019), disease mapping and risk (Zhang and Stram 2014; Atkinson et al. 2021), and identifying cases of genetic loci under selection (Tang et al. 2007; Zhou et al. 2016; Cuadros-Espinoza et al. 2022; Irving-Pease et al. 2024).

Although admixture is a common phenomenon across the tree of life, the majority of LAI methods have been developed and tested (reviewed in Geza et al. 2019; Schubert et al. 2020) with a focus on human genomics (Henn et al. 2012; Reich et al. 2012; Martiniano et al. 2017; Durvasula and Sankararaman 2019); a tendency largely driven by the greater availability of genomic data from humans compared to other species. Nevertheless, recent studies have showcased the importance of applying these methods in other species, primarily in animal breeds (Huson et al. 2012; Medugorac et al. 2017; Avallone et al. 2020; Chen et al. 2020; Donner et al. 2023), with a few providing alternative tools which can be applied to a range of other species (Dias-Alves et al. 2018; Oriol Sabat et al. 2022; Mora-Márquez et al. 2023).

Over the past few years, advances in laboratory protocols and computational tools have made it possible to retrieve hundreds of ancient genomes, from individuals that lived many thousands of years ago (Narasimhan et al. 2019; Allentoft et al. 2024). Additionally, the imputation of ancient human and non-human genomes (Martiniano et al. 2017; Erven et al. 2022, 2024, 2025; Todd et al. 2022, 2023; Sousa Da Mota et al. 2023; Allentoft et al. 2024; Bougiouri et al. 2025) has made it possible to reconstruct haplotypes with a high degree of accuracy, even in relatively low-coverage samples. This has enabled the ability to apply haplotype-based methods (such as LAI) to ancient genomes, something previously only possible with high-quality genomes from present-day samples. In principle, imputed ancient genomes could enable LAI using source proxies much closer in time to the admixture events under study, potentially leading to significantly higher accuracy. Indeed, a few ancient human studies have already made attempts to apply LAI methods on imputed samples from Mesolithic and Neolithic Eurasia (Martiniano et al. 2017; Haber et al. 2020; Pearson and Durbin 2023; Irving-Pease et al. 2024). However, detailed benchmarking and evaluation of the accuracy of LAI on ancient genomic data is currently lacking.

Despite the promising potential of LAI, there are several issues specific to ancient DNA (aDNA) which can affect the accuracy of inference. Poor DNA preservation, and other biases in sampling, can lead to small sample sizes and a distribution of data that is heterogeneous across both space and time. This may limit the number of samples that can be used as source populations for LAI in aDNA studies. Consequently, benchmarking LAI methods under various sampling schemes of present-day and ancient genomic data, particularly with respect to uneven temporal distributions of sampling, is of critical importance. To our knowledge, only one prior study has evaluated the impact of small source sample sizes on LAI accuracy with a focus on ancient datasets (Oliveira et al. 2024). However, the authors did so without considering time series data, thus missing a major feature of aDNA studies. Furthermore, as most LAI methods have been developed for scenarios involving recent admixture, thorough benchmarking of LAI accuracy as a function of admixture time is critical to establish the temporal boundaries for studying ancient admixture events. For example, the authors of FLARE benchmarked their approach simulating two, three and four-way models based on African, European and Asian demographic history, incorporating admixture happening only 12 generations ago (Browning et al. 2023), while MOSAIC simulated similar demographic models, extending the timing of admixture only back to 50 generations ago (Salter-Townshend and Myers 2019).

Here, we benchmarked a set of four popular LAI methods (FLARE, MOSAIC, RFMix, and simpLAI) (Maples et al. 2013; Salter-Townshend and Myers 2019; Browning et al. 2023; Oliveira et al. 2024) using extensive population genetic simulations, with a focus on sampling schemes common to ancient genomes. The insights we gained are broadly applicable to any organism with similar amounts and quality of genomic data. Utilizing the *slendr* simulation package, and its interface to the tree-sequence library *tskit* (Kelleher et al. 2018), we extracted the exact base-pair coordinates of true ancestry tracts in various simulation scenarios. We then inferred local ancestry tracts in the simulated genomes using four LAI methods. When applicable, we also compared the inferred admixture times with those inferred with DATES (Chintalapati et al. 2022), an alternative method for dating admixture events. By comparing the inferred ancestry tracts and admixture dates to the ground truth obtained from simulations, we evaluated the accuracy of LAI and admixture dating based on i) the timing of the admixture event, ii) the number of source individuals used and iii) the incorporation of non-contemporary sources, both prior to and after admixture, either when sampled from a single period or when grouped across periods. Our results serve as guidelines for potential LAI users, given the type of data available and research questions of interest.

## Methods

### Genetic simulations

We simulated genomic data in the form of tree-sequence files using the *slendr* R package (v.0.8) (Petr et al. 2023) under scenarios applicable to a range of real-world demographic histories. We tested a simple demographic scenario of one ancestral population (“population A”, *N_e_* = 10,000) splitting into two populations 1,500 generations ago (“population B” with *N_e_* = 10,000 and “population C” with *N_e_* = 10,000) (Fig. 1). These two populations were used as sources for the local ancestry inference. A new population (“population mix”, *N_e_* = 10,000) branches from one of the sources (population B, major ancestry) while also receiving gene-flow from the second source (population C, minor ancestry) at a rate of either 10% or 30% at a given number of generations ago (t_admix_). “Population mix” is the target admixed population, from which each haplotype in a given simulated genome is assigned an ancestry deriving from the two source populations. For each simulation we extracted 10 diploid samples from the target population at t_sampling_= 0 generations before present, and up to 100 samples from each of the two source populations at eleven different time points (see details of the sampling scheme below).

**Fig. 1:**
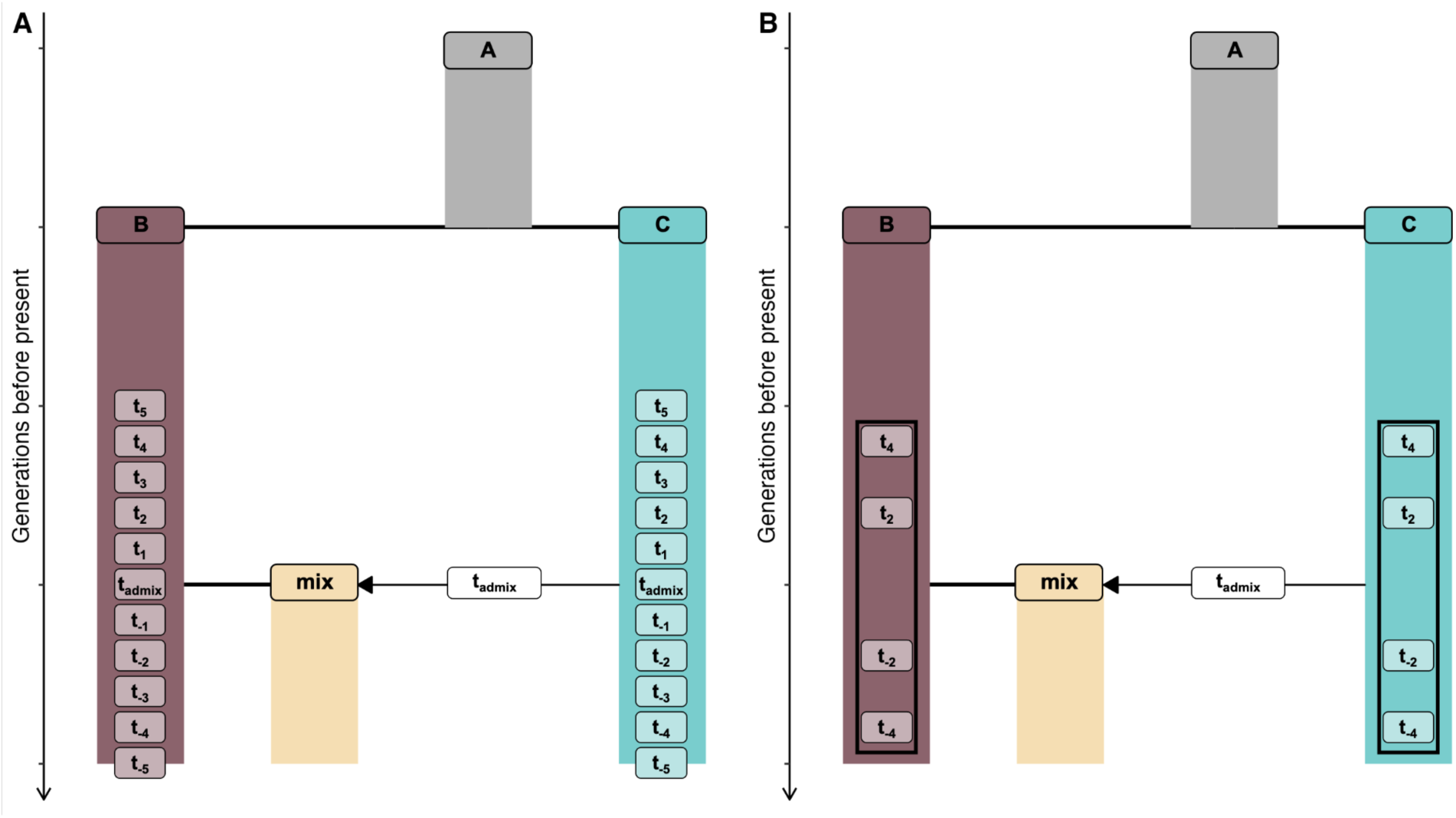
A schematic describing the simulated demographic models of gene-flow from two source populations (“B” and “C”) into the target population (“mix”), along with A) the timing of admixture in generations before the present (t_admix_=50, 100, 200, 300, 400 and 500) and time points at which the samples representing sources were recorded in a tree sequence, relative to the admixture time (t_sampling_ : t_-5_=0, t_-4_=t_admix_×0.2, t_-3_=t_admix_×0.4, t_-2_=t_admix_×0.6, t_-1_=t_admix_×0.8, t_0_=t_admix_, t_1_=t_admix_×1.2, t_2_=t_admix_×1.4, t_3_=t_admix_×1.6, t_4_=t_admix_×1.8, t_5_=t_admix_×2), and B) the four time points at which source samples were taken from and grouped for the time-series analysis (t_sampling_ : t_-4_=t_admix_×0.2, t_-2_=t_admix_×0.6, t_2_=t_admix_×1.4, t_4_=t_admix_×1.8).

For each demographic model, we simulated a tree-sequence using *slendr*’s coalescent simulation engine implemented in *msprime* (Baumdicker et al. 2022), and then converted it to a VCF format (Danecek et al. 2011) using the `ts_vcf()` function. For each simulation scenario, we simulated a sequence length of 300Mb, using a generation time of 1 (i.e., all of our results are represented in the time scale of generations), a uniform recombination rate of 10^-8^ per generation per base pair and a mutation rate of 10^-8^ mutations per generation per base pair. True ancestry tracts were extracted for each of the 10 target samples using the `ts_tracts()`function implemented within *slendr* based on the *tspop* framework (Tsambos et al. 2023), which returns the exact coordinates of ancestry tracts resulting from a given gene-flow event.

We built upon this model by modifying three different parameters and evaluating how they would affect the accuracy of downstream local ancestry inference:

● The number of samples taken from each of the two source populations (populations B and C) at n=2, 4, 6, 8, 10, 12, 14, 16, 20, 40, 50, 80 and 100.
● The timing of the admixture event between the two sources: t_admix_= 50, 100, 200, 300, 400 and 500 generations before present (Fig. 1A).
● The time point at which the source samples were taken from, relative to the admixture time, t_sampling_: t_-5_=0, t_-4_=t_admix_×0.2, t_-3_=t_admix_×0.4, t_-2_=t_admix_×0.6, t_-1_=t_admix_×0.8, t_0_=t_admix_, t_1_=t_admix_×1.2, t_2_=t_admix_×1.4, t_3_=t_admix_×1.6, t_4_=t_admix_×1.8, t_5_=t_admix_×2 (Fig. 1A). We also tested whether using a merged time-series dataset of source samples from multiple time periods would affect the results. We did this using four selected time points pre- and post-dating the admixture event (t_-4_=t_admix_×0.2, t_-2_=t_admix_×0.6, t_2_=t_admix_×1.4, and t_4_=t_admix_×1.8). From each of these four time points we extracted either 1, 2, 3 or 4 individuals, adding up to a population size of n=4, 8, 12 or 16 from each of the two sources (Fig. 1B).

To capture the effect of stochastic variation in the simulated data, we used 10 random seeds to simulate 10 replicates of each demographic model. We computed F_ST_ between the two source populations using the *slendr* function `ts_fst()`.

### Local ancestry inference

We filtered each VCF with the genotypes of all simulated individuals for biallelic sites and applied a minor allele frequency (MAF) cutoff of 0.01. Each simulated VCF contained phased haplotypes and was used as input for four local ancestry inference methods evaluated in this study: FLARE (Browning et al. 2023), MOSAIC (Salter-Townshend and Myers 2019), RFMix (Maples et al. 2013) and simpLAI (Oliveira et al. 2024). All parameters were left to default for each method, unless specified below.

FLARE has been shown to achieve high accuracy with reduced memory requirements and computation time compared to other methods, allowing LAI in large reference panels. We ran FLARE v0.3.0 specifying the 10 target samples as one population to infer local ancestry, and the samples from populations B and C as the two source populations. For each simulated VCF, we specified the exact time of admixture (t_admix_) used for that simulation as the `gen` command-line parameter of FLARE. We also specified a `min-mac` of 1 and enabled reporting of the posterior ancestry probabilities using the `prob=true` flag.

MOSAIC permits an arbitrary number of source ancestries, and has shown high accuracy when using small reference panel sizes (Browning et al. 2023). As input, we prepared three different haplotype files containing either the target samples, the samples from source population B or the ones from source population C. Following Browning et al. 2023, we specified the number of grid points per cM (`--GpcM`) to be the product of 0.0012 and the number of sites within the simulated VCF, as this was shown to improve the LAI of MOSAIC for sequence data in their study. We ran MOSAIC v1.5.0 specifying 10 target individuals (`--number 10`), two ancestries (`--ancestries 2`) and the exact time of admixture used for that simulation (t_admix_) as the `--gens` parameter. Given that the correct phase is known from our simulations, we enabled the `--nophase` option to prevent MOSAIC from attempting to correct potential phasing errors.

RFMix is a popular machine learning LAI tool which relies on random forests with a conditional random field (Maples et al. 2013). We ran RFMix v2.03-r0 and as input we provided two VCF files, one with the 10 target samples, and one with the samples from source populations B and C. As RFMix requires an input admixture time, we specified here the exact time of the admixture used for the simulations (t_admix_) as the `-G` parameter. We used the `--reanalyze-reference` option to iteratively analyze the reference haplotypes in addition to the target haplotypes, and enabled the expectation-maximization (EM) optimization option (`-e 5`) to perform a maximum of 5 iterations.

simpLAI is a recently developed method tailored for LAI from small reference panels (Oliveira et al. 2024). As input, we used a genotype file containing samples from the target and source populations and then specifying the number of total haplotypes present within each (`--ssa, --ss1, --ss2`). We ran simpLAI v0.9 using the *rec* mode, which accounts for intra-population ancestry switches that may have happened within a haplotype through recombination. For this mode, we specified 2000 polymorphic sites to search for the shortest recombination path (`-n 2000`), as recommended by the developers for older admixture times, and kept the defaults of 1000 sites as an increment for the sliding window when searching for the shortest recombination path (`-m 1000`), and 5 linked polymorphic sites between potential recombination breakpoints (`-t 5`).

For the evaluation of FLARE, MOSAIC and RFMix on simulated data, we used a recombination map with a uniform rate of 10^-6^ cM/Mb (simpLAI does not require a recombination map). The output of all four LAI methods contains the ancestry assignment for each of the two haplotypes of each site in the VCF, for which a posterior probability is also available in the outputs from FLARE, MOSAIC and RFMix. Given that posterior probabilities are not equally calibrated across different methods, we first explored their distribution from each method’s output, when applicable. Because the purpose of our downstream anayses is on the accuracy of inferred ancestry tracts, we converted the local ancestry of each individual site to continuous segments by assigning the ancestry of each individual site not present in the simulated dataset, to that of the closest preceding site. The ranges of each ancestry tract were then determined as runs of consecutive sites assigned to ancestry 1 or ancestry 2.

### Overlap estimation and accuracy metrics

We calculated the per base-pair overlap of the inferred ancestry tracts from each of the four methods in relation to the true tracts using the GenomicRanges R package (v1.50.2) (Lawrence et al. 2013). We estimated the true positives (TP), false positives (FP), true negatives (TN) and false negatives (FN) for ancestry B (Fig. S1), noting that the quantities for ancestry C are the exact complements (for instance, TP for ancestry B automatically implies TN for ancestry C). We then computed the Matthews correlation coefficient (MCC) (Matthews 1975) as a metric to evaluate the overall accuracy of the local ancestry inference. The MCC metric is calculated based on all four confusion matrix categories (FN, FP, TN, TP):

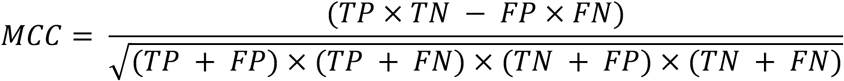

A high MCC score is achieved when all four confusion matrix metrics yield high true rates and low false rates, rewarding both correct positive and correct negative predictions. To calculate the MCC, we used the `mcc` function from the *mltools* R package (Ben Gorman 2018) and then calculated a normalized version of the MCC, called nMCC (Chicco and Jurman 2020):

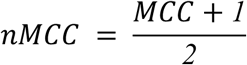

nMCC ranges from 0 to 1, where a value equal to 0.5 indicates an arbitrary prediction and a value closer to 1 represents high overlap with the truth. nMCC estimates were averaged across the 10 target samples for each run.

In order to aid the interpretation of the patterns produced by the nMCC metric and its ability to weigh all elements of a confusion matrix together, we also calculated sensitivity (true positive rate) for both ancestry B and C:

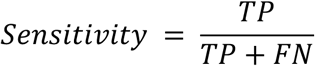

### Admixture time inference

Among the four LAI methods tested, two (FLARE and MOSAIC) also provide estimates of admixture dates. FLARE outputs an inferred number of generations since admixture. To infer admixture dates, MOSAIC relies on the co-ancestry curves from (Hellenthal et al. 2014), by fitting the exponential decay of the ratio of probabilities of pairs of local ancestries as a function of genetic distance. Each of these curves depends on an admixture parameter lambda (λ), which governs the exponential decay of tract lengths since admixture. For our given two-way admixture model, we averaged across the 3 estimates of the λ parameter outputted by MOSAIC, in order to report an admixture time.

In order to compare the accuracy of the haplotype-based admixture dating, we additionally tested DATES, a commonly used admixture dating tool tailored for low coverage ancient genomes that does not require phased data (Chintalapati et al. 2022). For DATES, we converted the simulated genomic data containing the target and source populations into eigenstrat format using CONVERTF (https://github.com/argriffing/eigensoft/tree/master/CONVERTF). We then ran DATES v4010 specifying the recommended bin size of 0.001 Morgans (`binsize: 0.001`), a maximum distance of 1 Morgans (`maxdis: 1`), a random seed (`seed: 77`), and enabled the exponential fit (`runfit: YES`) and the use of affine for the fit (`afffit: YES`). We note that the calculation of confidence intervals on simulations (which DATES computes by jackknifing across multiple chromosomes) was disabled (`jackknife: NO`) as our simulated data consists of only one chromosome. Instead, to present the accuracy of DATES inference in the context of the inherent stochasticity in the data, we performed our analyses across the 10 replicates of each simulated scenario. Furthermore, since we used a recombination map with a uniform rate, the genetic and physical positions in the simulated data are highly correlated. Therefore, we opted for the `checkmap: NO` option.

### Admixture proportions

Admixture proportions were estimated by summing up the total length of tracts assigned to each of the two ancestries, divided by the total genome sequence length. We further tested the impact of applying posterior probability cutoffs on the total sequence length retained, as well as the inferred admixture proportions estimated.

## Results

We assessed the performance of the four chosen LAI methods, FLARE, MOSAIC, RFMix and simpLAI, on demographic models (Fig. 1) simulated across all combinations of three parameters: the admixture time (t_admix_), the number of samples taken from each source population (n) and the time points from which the source individuals were sampled from (t_sampling_), intended to capture a broad range of modeling situations in practice. We then quantified the degree of overlap between the inferred tracts and the true tracts known from the simulations, as well as the difference between the true and inferred admixture times and proportions.

### Accuracy of inferred local ancestry tracts

We examined the overlap of the inferred tracts from MOSAIC, FLARE, RFMix and simpLAI with the true tracts for each of the two source ancestries, populations B and C. For each simulated scenario, we calculated the total amount of sequence as TP, FP, TN and FN in base pairs, and used them to compute the i) nMCC accuracy metric (Fig. 2, Fig. S2), and ii) sensitivity (Fig. S3, S4).

**Fig. 2:**
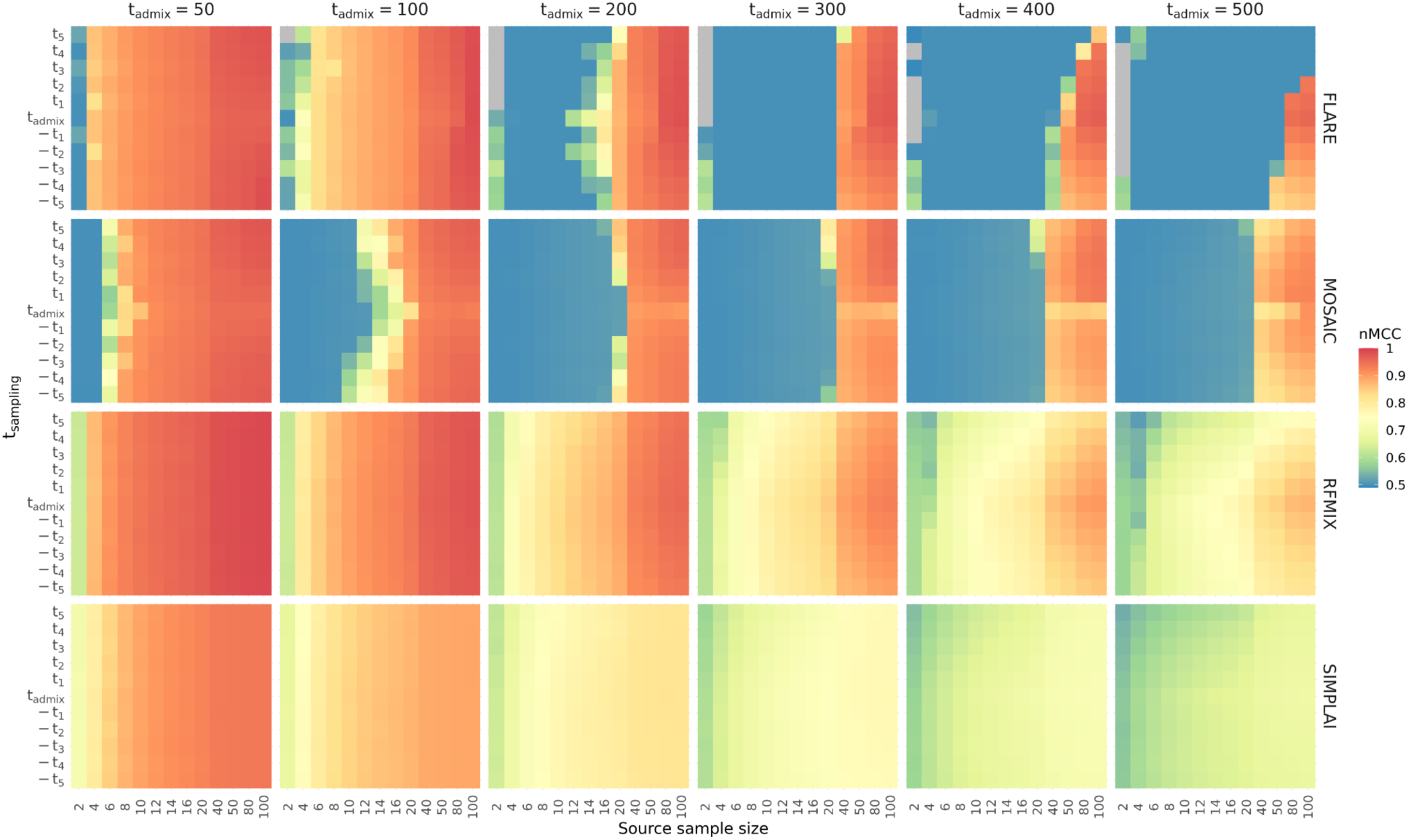
nMCC values quantifying the overlap of the inferred and true tracts for the four methods. Due to the complementary nature of all four confusion matrix categories (TP, FP, TN, FN), the nMCC values are the same for the tracts assigned to either of the two ancestries. The y-axis represents the 11 time points at which the source samples were taken from (t_sampling_), relative to the admixture event time (t_admix_). The x-axis is the number of samples taken from each source population. Each column represents the six tested admixture times (t_admix_), using a gene-flow rate of 30%, and each row the results for the four methods. Each square shows the average across all 10 samples from the admixed population and across all 10 replicates. An nMCC value equal to 0.5 (blue) indicates an arbitrary prediction and a value closer to 1 (red) represents high overlap. Grey indicates cases where at least one individual from the target population was inferred to have ancestry tracts that did not match any true tracts (TP = 0, FP ≠ 0).

The nMCC metric is calculated using all four estimates: TP, FP, TN, FN, and therefore is the same when focusing on tracts assigned to population B or population C (for instance, a TP of one ancestry necessarily implies a TN of the other ancestry, Fig. S1). As expected, all four methods showed high nMCC scores (> 0.8) for scenarios with recent admixture times and lower sample sizes, with LAI for older admixture events accurate only with sufficiently large source sample sizes (Fig. 2). This pattern is consistent for both gene-flow rates tested (Fig. S2). Comparing the four methods, for given admixture times and small source sample sizes, FLARE and MOSAIC showed nMCC values close to 0.5 (shown by blue), effectively equivalent to an arbitrary assignment of ancestries, whereas RFMix and simpLAI showed a more gradual increase in accuracy with increasing source sample sizes across admixture times. For example, for t_admix_ = 50, FLARE and MOSAIC showed high nMCC estimates above 0.5 only when either four or six samples from each source were used respectively, whereas RFMix and simpLAI showed higher nMCC values using as few as 2 samples. When focusing on older admixture times (200 ≤ t_admix_ ≤ 300), FLARE and MOSAIC were not able to infer local ancestry (nMCC ≈ 0.5) until at least 40 samples were included from each source, whereas RFMix and simpLAI managed to infer ancestry tracts correctly with fewer samples, though with relatively low nMCC scores (nMCC < 0.8). When including large sample sizes (n ≥ 40) FLARE, MOSAIC and RFMix outperformed simpLAI, with nMCC scores above 0.8, even when approaching the oldest admixture times evaluated (400 ≤ t_admix_ ≤ 500). With lower gene-flow rates (10%), MOSAIC struggled to infer accurate ancestry tracts across all admixture times, unless at least 40 individuals per source were available (Fig. S2). FLARE showed a similar behaviour for t_admix_ = 200, however, struggled to recover accurate ancestry tracts for admixture times t_admix_ ≥ 400. RFMix and simpLAI showed similar performance under both tested gene-flow rates.

Focusing on the different source sampling times, there was no distinctive impact on accuracy regardless of whether the source proxies were sampled before or after the admixture event. For older admixture times (400 ≤ t_admix_ ≤ 500) and large source sample sizes, FLARE showed decreased accuracy when sources were sampled prior to the admixture event compared to after (Fig. 2, Fig. S2), a pattern which can be attributed to the low levels of differentiation between the two sources at these time points (F_ST_ < 0.02) (Fig. S5). Cases with low nMCC values seem to be driven by low sensitivity estimates, thus low true positive rates, for tracts assigned to the minor ancestry (population C) (Fig. S3, S4). This suggests that when source sample sizes are low, especially for older admixture times, FLARE and MOSAIC will tend to incorrectly infer the major ancestry in regions with minor ancestry.

When comparing the true and inferred tract lengths (Fig. S6, S7), RFMix showed a consistent overlap of tract length distribution with the true tracts across demographic scenarios. MOSAIC inferred shorter tract lengths for both ancestries across admixture times, unless large source sample sizes were used (e.g. n = 20 for 30% gene-flow rate). In cases with low sample sizes or old admixture events, FLARE would only infer tracts assigned to the major ancestry, whereas larger sample sizes would result in a more accurate overlap of the tracts length distribution with that of the true tracts (Fig. S6, S7). Finally, simpLAI showed bimodal tract length distributions, primarily for the minor ancestry tracts, and tended to infer tracts longer than the true ones in both ancestries at older admixture times.

### Filtering for highly confident inferred tracks

Given that posterior probabilities produced by the benchmarked methods are not calibrated in a comparable, consistent way (or not at all, as is the case for simpLAI), we next evaluated whether applying posterior probability cutoffs would affect the total genome length retained and accuracy of the inferred tracts as well as the admixture proportions from the two source populations.

MOSAIC showed a decreasing amount of sequence retained with higher probability cutoffs across demographic scenarios, reflecting the broad probability distribution in Fig. S8, S9. However, for older admixture events or low sample sizes, the retained ancestry tracts showed nMCC scores below 0.8, even when > 75% of the sequence was kept (Fig. 3). This suggests that high posterior probability cutoffs do not necessarily reflect retaining accurate ancestry tracts in MOSAIC. For FLARE, for admixture times t_admix_ ≤ 100 generations before present or sample sizes n > 20, applying a posterior probability cutoff of 0.9 would retain more than 75% of the sequence with nMCC values above 0.8. In cases with older admixture events and lower sample sizes, 100% of the sequence was retained, regardless of the probability cutoff applied, though with nMCC values below 0.8. These results show that FLARE tends to incorrectly assign the entire haplotype to the major ancestry with high posterior probability for older admixture events or when low source sample sizes are present. Finally, applying posterior probability cutoffs on the tracts inferred by RFMix did not result in a change in the amount of sequence retained, in cases where nMCC values were either above or below 0.8. For lower gene-flow rates (10%), MOSAIC presented similar percentages of sequence retained, though with lower nMCC scores, while FLARE showed the same pattern of assigning the entire chromosome to a single ancestry at more recent admixture times and higher source sample sizes (Fig. S10). Overall, these results suggest that applying universal posterior probability cutoffs across LAI methods does not guarantee similar output and levels of accuracy, and requires manual inspection.

**Fig. 3:**
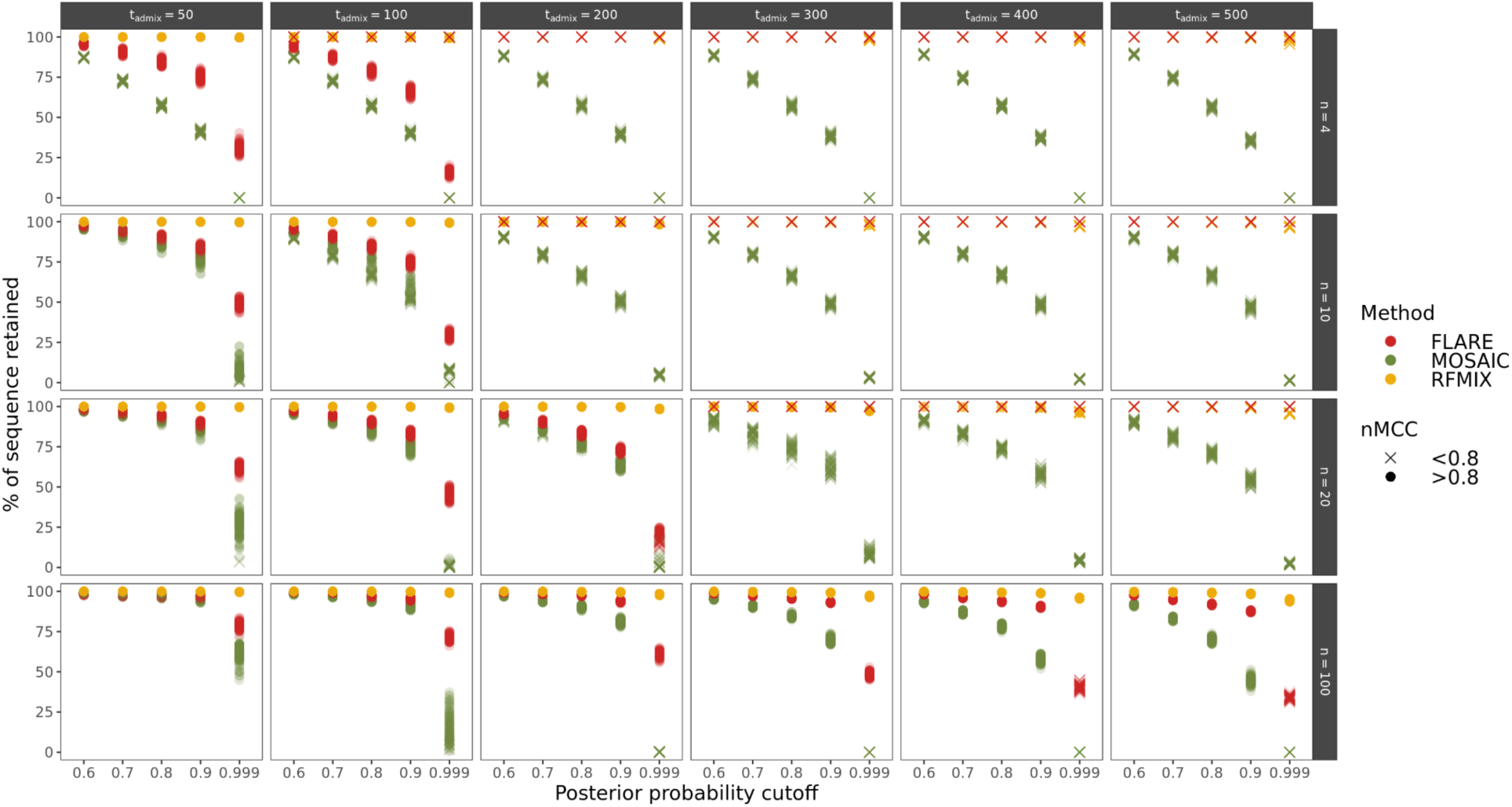
Proportion of sequence left after applying five different probability cutoffs. Each column represents the five tested admixture times (t_admix_), using a gene-flow rate of 30%, and each row a subset of four source sample sizes (n = 4,10,20,100). Results are shown for one source sampling time (t_sampling_ = 0 generations before present). The colour represents the method used and shape depicts nMCC values either above or below 0.8. Each point shows the estimated value for each of the 10 target samples from each of the 10 replicates. We note that simpLAI is missing in this comparison because it reports ancestry tracts for the entire genome without any posterior probability output.

We further explored the impact of posterior probability cutoffs on admixture proportion estimates from each method. When applying no cutoffs, all four methods inferred admixture proportions of the minor ancestry approximately equal to the true simulated values, when focusing on higher gene-flow rates (30%), recent admixture times and large source sample sizes (Fig. S11). Testing older admixture times or smaller sample sizes would result in admixture proportions of the minor ancestry reaching zero in FLARE and MOSAIC, whereas simpLAI and RFMix would show consistent estimates approximate to the true admixture proportions, even in cases where nMCC < 0.8. Applying posterior probability cutoffs on the three methods (since simpLAI does not report posterior probabilities) resulted in decreasing admixture proportion estimates from FLARE and MOSAIC, whereas RFMix produced consistent estimates around the true admixture proportion. Focusing on simulated scenarios with smaller gene-flow rates (10%), we observed similar results (Fig. S12). However, RFMix showed consistent overestimation of the minor ancestry admixture proportions across admixture times, sample sizes and posterior probability cutoffs (Fig. S12). These results reflect the percentage of inferred tracts retained from each method after applying posterior probability cutoffs, suggesting that applying them might bias admixture proportion estimates.

### Inferring admixture time

In addition to LAI, results of the FLARE and MOSAIC inference include an estimate of the timing of admixture events, expressed as “generations before the target”, via a single point estimate derived from all samples in the target population. When comparing this estimate to DATES–an allele frequency-based admixture dating method–both LAI methods were outperformed in inferring the admixture time across all demographic scenarios (Fig. 4, S13). In the range of 50 ≤ t_admix_ ≤ 100, MOSAIC was able to approximately infer the true admixture time across sample sizes and sampling time points, however, it showed large deviations at smaller sample sizes (n ≤ 8) for t_admix_ = 100. At older admixture times, MOSAIC consistently underestimated the admixture time across all source sample sizes. Overall, FLARE tended to overestimate the admixture time, reaching estimates more than twice the actual true value across all demographic scenarios. The same results were present with lower gene-flow rates (10%), with FLARE showing increased overestimations even at more recent time scales (Fig. S14).

**Fig. 4:**
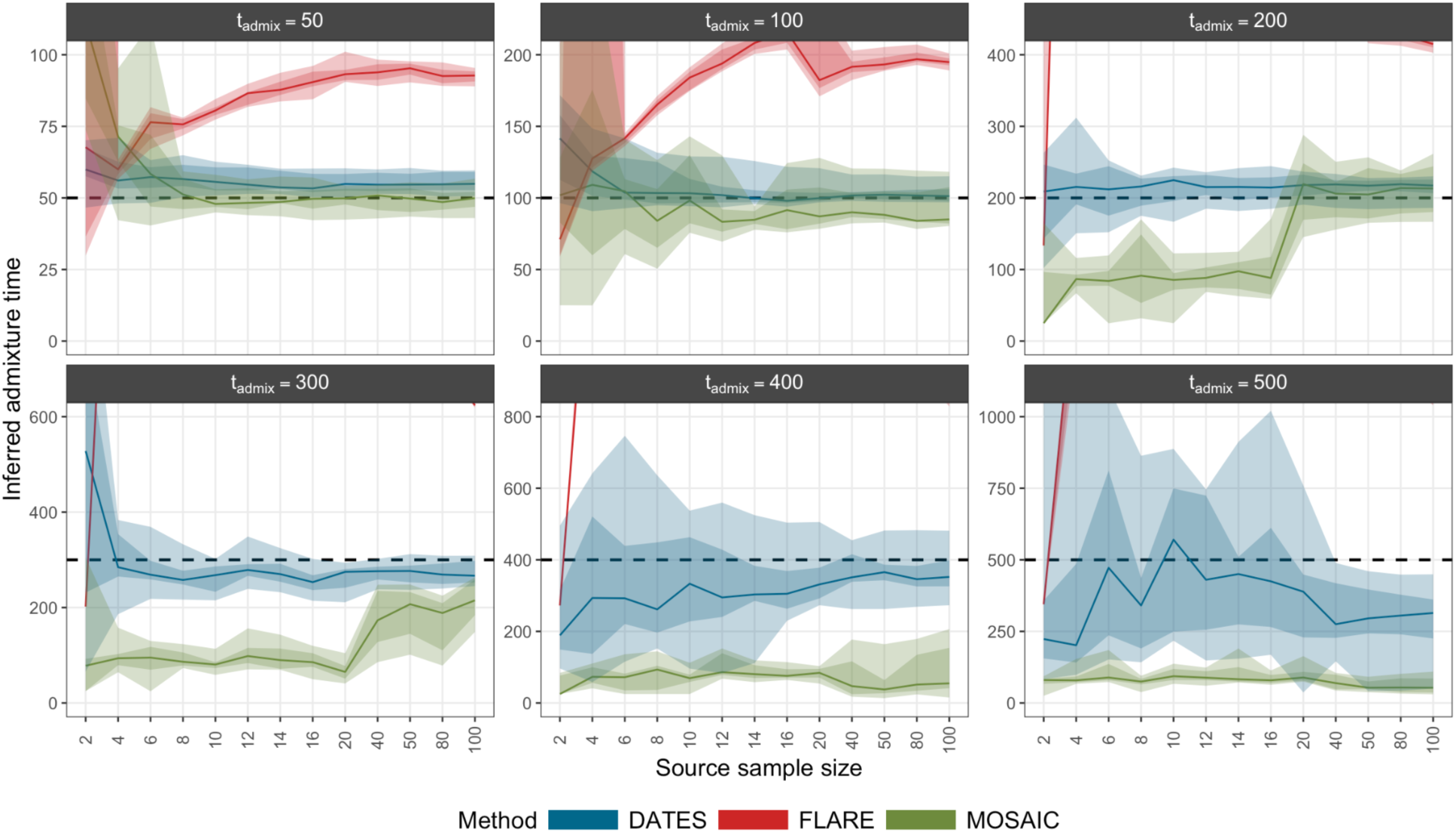
Inferred admixture times (y axis) under a subset of demographic parameters from DATES (blue) FLARE (red) MOSAIC (green). The x-axis shows the 13 sample sizes taken from each of the two ancestries. Each facet shows six different admixture times (t_admix_ = 50,100,200,300,400,500) using the same source sampling time (t_sampling_ = 0 generations before present) and a gene-flow rate of 30%. The dotted black line indicates the true admixture time. The shaded regions in each ribbon show the 50% and 80% quantiles and the line represents the median. Estimates which fall outside of the range [0, 2×t_admix_] are not shown. All combinations of demographic parameters are shown in Fig. S13, S14.

DATES produced accurate estimates for admixture times up to t_admix_ = 300 generations prior to the target population, with a tendency to slightly overestimate at more recent admixture times (t_admix_ ≤ 200). Going towards even older events (t_admix_ ≥ 400), inferred admixture times were underestimated, with minor changes observed with larger sample sizes. In scenarios where the source sample size was quite low (n = 2), DATES struggled to correctly infer the admixture time, even at recent admixture time (t_admix_ ≤ 100). Chintalapati et al. (2022) suggested that estimated admixture times as inferred from DATES should be considered significant if the admixture event was less than 200 generations old and the normalized root-mean-square deviation (NRMSD) was < 0.7. Thus, we assessed whether the NRMSD scores could serve as a reliable indicator for the accurate inference of admixture times. Overall, in our tested scenarios where DATES showed accurate results for t_admix_ ≤ 300, NRMSD values were < 0.4 (Fig. S15, S16). For t_admix_ = 400, where DATES underestimated the admixture time, NRMSD values exceeded 0.7, primarily for sample sizes below 10, whereas for t_admix_ = 500, cases where NRMSD estimates exceeded 0.7 were observed across all sample sizes. However, we note that there were cases in which DATES would report NRMSD values < 0.7, but severely misestimate the admixture times, primarily at low sample sizes. In cases with lower gene-flow rates (10%), DATES was accurate only at more recent time scales (t_admix_ ≤ 200) and larger sample sizes.

### Time series dataset

In addition to testing the effect of sampling source individuals from different time points on LAI accuracy, we also explored the impact of combining sources from different time points (Fig. 1B). This is particularly relevant because the scarcity of aDNA sampling means that, in practice, a source proxy panel is rarely composed of individuals sampled from the same exact point in space and time. We selected samples across four distinct time points relative to the admixture event (t_-4_ = t_admix_×0.2, t_-2_ = t_admix_×0.6, t_2_ = t_admix_×1.4, and t_4_ = t_admix_×1.8). These corresponded to grouping of samples which were sampled between 10 and 90 generations ago for t_admix_ = 50, between 20 and 180 generations ago for t_admix_ = 100, between 40 and 360 generations ago for t_admix_ = 200, between 60 and 540 generations ago for t_admix_ = 300, between 80 and 720 generations ago for t_admix_ = 400 and between 100 and 900 generations ago for t_admix_ = 500. The total number of samples from each source amounted to 4, 8, 12, or 16 individuals. We observed that the metrics of accuracy between inferred and true ancestry tracts remained stable, showing neither improvement nor deterioration, irrespective of source sample size, admixture time and gene-flow rates (Fig. S17-S20). Additionally, the estimated admixture times were consistent, regardless of whether the data was analyzed as a time series or by sampling from a single time period (Fig. S21, S22). This suggests that under demographic scenarios such as those tested here, with population continuity post admixture, grouping samples from different time points into one source population does not impact the overall LAI results and ancestry source proxies can be safely combined from samples from multiple time snapshots.

### Example scenarios

Finally, we demonstrated the performance of all methods on a pair of contrasting example scenarios featuring an admixture event occurring either 50 or 200 generations ago, both with a gene-flow rate of 30% and using 10 samples per source population sampled at the present (t_sampling_ = 0) (Fig. 5). In each case, the average tract length of the minor ancestry is 2.7 Mb and 0.68 Mb respectively. For the recent admixture event, all methods showed a high overlap of inferred ancestry tracts with the true tracts known from the simulations (Fig. 5Ai), with the inferred tract length densities matching the true densities and high posterior probabilities for assigning either ancestry (Fig. 5Aii,iii). nMCC scores were above 0.8 across methods, and applying posterior probability cutoffs did not affect accuracy levels, however, it reduced the overall percentage of sequence retained in FLARE and MOSAIC (Fig. 5iv). Finally, the inferred admixture time from DATES was 55.3 generations ago, and closely matched the true admixture time, with a low NRMSD value of 0.047 (Fig. 5Av). For the older admixture event, RFMix showed better performance in recovering the true tracks compared to the other three methods (Fig. 5Bi), with tract length density similar to the true one and overall high posterior probabilities and nMCC rates (Fig. 5Bii, iii, iv). simpLAI overestimated the overall tract length, whereas MOSAIC underestimated it and showed low nMCC scores across all posterior probability cutoffs (Fig. 5Bii,iv). FLARE inferred the whole genome as the major ancestry with high posterior probability (Fig. 5Bi, iii). DATES inferred an admixture time around 248.5 generations ago (Fig. 5Bv), aligning again with the true admixture time of 200 generations. Focusing on the overall nMCC estimated across the simulated scenarios in Fig. 2, we see that for the scenario shown in Fig. 5B, FLARE and MOSAIC nMCC values were at 0.5, suggesting a complete random ancestry assignment, whereas RFMix and simpLAI showed estimates around 0.8.

**Fig. 5:**
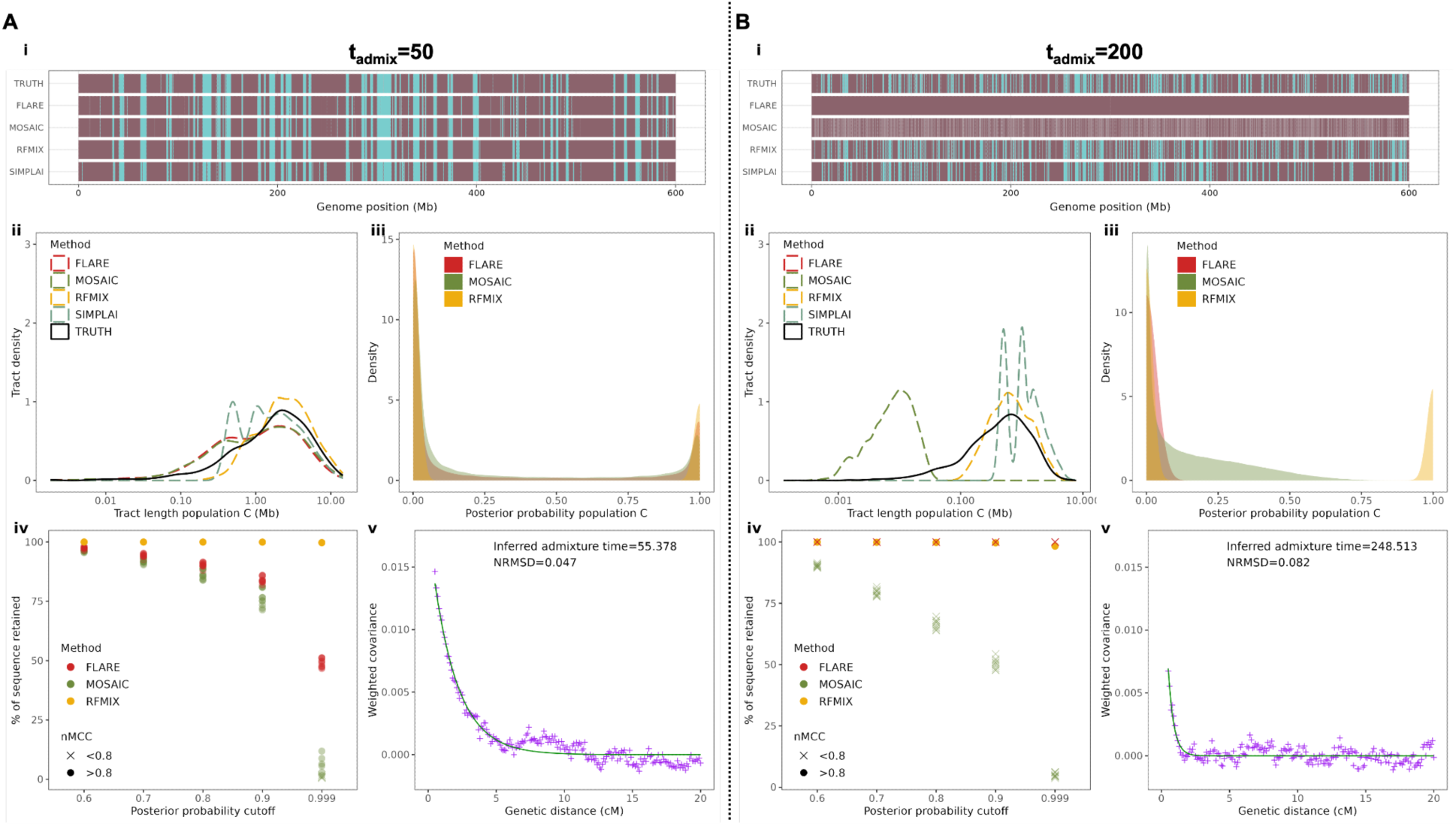
Two example demographic scenarios for a recent (t_admix_ = 50 generations ago) and older (t_admix_= 200 generations ago), using a gene-flow rate of 30% and the LAI output from the four tested methods. A) t_admix_= 50 generations ago, n = 10 samples per source, t_sampling_ = 0 generations before present from a single simulation run. B) t_admix_= 200 generations ago, n = 10 samples per source, t_sampling_ = 0 generations before present from a single simulation run. i) True and inferred tracts for one individual, with purple representing population B and blue population C. ii) Tract length density plot for true and inferred population C tracts. iii) Posterior probability density plot for inferred tracts assigned to population C. iv) Percentage of sequence left after applying five different posterior probability cutoffs. The colour represents the method used and shape depicts nMCC values either above or below 0.8. Each point shows the estimated value across all 10 target samples. We note that simpLAI is missing in figures iii and iv because it reports ancestry tracts for the entire genome without any probability filtering. v) DATES ancestry covariance curve with the reported admixture time and NRMSD estimation.

## Discussion

In this study, we investigated the accuracy and potential of LAI for ancient DNA studies, particulary in the context of the inherent sampling heterogeneity, focusing on four published LAI methods FLARE, MOSAIC, RFMix, and simpLAI. We performed an extensive set of simulation benchmarks under a variety of demographic scenarios capturing a broad range of practical situations, and evaluated how well these infer the coordinates of local ancestry tracts and estimate admixture times and proportions. To measure the inference accuracy of all methods, we leveraged the tree sequence data structure as a means to precisely and efficiently identify the coordinates of true ancestral tracts within each simulation scenario.

In agreement with previous findings (Browning et al. 2023), we found that all methods achieved higher accuracy metrics with increasing sample sizes of the proxy source populations. The accuracy of inference with smaller sample sizes was highly dependent on the timing of the admixture event, with larger sample sizes required for accurate inferences for older admixture events. FLARE and MOSAIC showed a striking dependence of LAI accuracy on the source sample size, whereas RFMix and simpLAI showed more robustness to small sample sizes. In general, only a few studies have addressed the number of samples used as sources as a crucial factor in LAI accuracy (Baran et al. 2012; Maples et al. 2013; Durvasula and Sankararaman 2019), with some only considering memory requirements and runtime (Schubert et al. 2020). However, only one study has tested inference for sample sizes as low as those present in this study (2 ≤ n ≤10) which are much more practically relevant for ancient genomic studies (Oliveira et al. 2024). Our results showed that for recent admixture times, RFMix and simpLAI achieved high results for low samples sizes, where simpLAI showed good performance using as few as two samples per source population for an admixture time of 50 generations, demonstrating its utility as a useful tool for low sample sizes and recent admixture events (Oliveira et al. 2024). In cases with larger sample sizes with at least 40 samples per source, FLARE, MOSAIC and RFMix performed well for older admixture times, up to 500 generations. We note that lower gene-flow rates would require more individuals per source population to reach accuracy levels comparable to scenarios involving higher gene-flow rates, with MOSAIC and FLARE displaying the largest changes in accuracy between the two scenarios (Fig. 2, S2).

Filtering for ancestry tracks based on the posterior probability of ancestral assignment as a confidence criterion produced varying outcomes, depending on the method and demographic parameters tested. In the most challenging cases, involving old admixture times or low sample sizes, FLARE had a tendency to incorrectly assign the entire genome with high confidence to the major ancestry. MOSAIC showed a pattern of decreasing total sequence length retained after the probability cutoff filtering, which would often not result in higher accuracy. RFMix was more robust when applying such probability filters, with large amounts of sequence retained even at high probability cutoffs (> 0.9), without influencing inferred ancestry proportions in a significant way (Fig. S11). In contrast, ancestry tracts assigned to the minor ancestry were filtered out at higher probability cutoffs in MOSAIC and FLARE (apart from cases where the whole genome was assigned the major ancestry), thus impacting the overall estimated admixture proportions. Overall, no universal rule can be established for filtering based on a posterior probability cutoff of ancestry assignments across all scenarios. The choice of such cutoff should be method dependent, and include an initial inspection of the overall posterior probability distribution. For example, RFMix presented a bimodal distribution of posterior probabilities for the minor ancestry, whereas MOSAIC had a more uniform distribution extended across the whole posterior probability range (Fig. S8, S9). Consequently, applying a given posterior probability cutoff would remove tracts in one case, but not the other. A successful inference requires careful consideration regarding the amount of sequence retained after filtering and the admixture proportions estimated.

Crucially, our analyses suggest that admixture events investigated using LAI can be pushed back towards much older time scales than those investigated in the original publications of the benchmarked methods (Maples et al. 2013; Salter-Townshend and Myers 2019; Browning et al. 2023; Oliveira et al. 2024), all of which have been developed and tested for admixture events for more recent time scales, generally less than 50 generations. The majority of LAI methods and studies have focused on admixture events during recent human history, such as those that occurred in African-American (Jin et al. 2012; Maples et al. 2013; Schubert et al. 2020), Northern Native American (Reich et al. 2012; Maples et al. 2013) or Latin American populations (Baran et al. 2012; Deng et al. 2016). Our analyses suggest that for an admixture event that occurred 500 generations ago, accurate inferences would require a sample size of at least 40 individuals, given a recombination rate of 10^-8^ per generation per base pair. Given this recombination rate, and using generation times of species such as humans, cattle or dogs, our results would translate to inferring local ancestry for admixture events which happened 10,000, 3,000 or 1,500 years before the sampling of the target population, respectively. Given the parameter ranges we investigated and the increasing amount of ancient human and animal domesticate data generated across these time-scales (Bergström et al. 2020; Allentoft et al. 2024; Cai et al. 2025), this would mean that inferring ancestry tracts in Bronze Age, Iron Age and Neolithic genomes can be quite accurately done, depending on the generation time of the species. Therefore, the conclusions and lessons learned can readily be applied as a guideline for different species. We note, however, that inferring local ancestry at very old admixture times would have its own natural upper limits, which would also be dependent on the recombination rate. As we move towards older admixture times, tracts would tend to become shorter and shorter, up to a point where their inference would be impossible, with higher recombination rates speeding up this process. Under the demographic scenarios and recombination rate considered here, the shortest average tract length for the minor ancestry in samples descending from an admixture event 500 generations ago with gene-flow rates of 30% and 10% was 0.27 and 0.22 Mb respectively (Fig. S6, S7).

A major prerequisite to navigate the parameter grid in Figure 2 is having an *a priori* idea of how old the admixture event is relative to the target population, in order to see whether it fits within our suggested time limits for LAI. Furthermore, somewhat counterintuitively, many LAI methods require a point estimate admixture time as input parameter. This is usually unknown, and assuming an incorrect time of admixture might bias the LAI outcome. Oliveira et al. (2024) suggested that for methods such as RFMix, carrying out inferences by testing different input admixture times and filtering for genomic regions for which the LAI is consistent might be one approach to deal with such uncertainties. However, we show that filtering out regions might, in fact, bias the estimated admixture proportions (Fig. S11, S12). To circumvent this issue, we investigated the possibility of using the DATES admixture dating method as a more robust first step, in order to then provide an approximate estimate of how far back the admixture event could be to an LAI method. Based on our simulations, DATES was able to accurately infer the admixture time for events as old as 300 generations prior to the sampling time of the target population, given adequate sample sizes. Furthermore, DATES outperformed the two tested LAI methods (FLARE and MOSAIC) which also provide admixture time estimates, even when they were given the exact admixture time as input. In order to assess whether the results from DATES were reliable, we referred to the NRMSD values produced by the software (Chintalapati et al. 2022). We observed that accurately inferred admixture dates had lower NRMSD rates (< 0.2), whereas higher values were often associated with misestimated admixture times, even when they remained under the suggested NRMSD cutoff of 0.7 (Chintalapati et al. 2022) (Fig. S15, S16). Therefore, we suggest using DATES as a complementary investigation step prior to LAI to establish an approximate admixture time, while applying a stricter NRMSD cutoff, especially for older admixture times. In fact, since the LAI methods tested here seem to struggle with precise admixture time inference in a wide range of simulated scenarios suggests that DATES should perhaps be considered as a primary inference method for this purpose.

Grouping individuals from various time points into a single source population did not adversely affect the accuracy of LAI. This finding is particularly relevant when present-day individuals do not adequately represent proxies as sources for older admixture events, or when data is unevenly distributed across a time series transect—a scenario often encountered in studies of aDNA. Overall, grouping present-day, historical or ancient samples into joint source reference panels could increase the effective sample sizes needed for precise determination of local ancestry tracts and admixture timings. However, this approach presupposes an understanding of the source population’s continuity, as past admixture events may have influenced some source samples differently than others. Further assessment of the impact of admixture events occurring throughout the source populations’ evolutionary histories will be important to address the extent to which this would impact LAI accuracy.

We note that although we focused only on the performance of four LAI methods, future studies will be able to easily build upon our reproducible benchmarking pipeline by evaluating the performance of other tools which might potentially perform better in scenarios where the four methods tested here performed poorly. Investigating the behavior of tools such as Loter (Dias-Alves et al. 2018), ELAI (Guan 2014) and LAMP-LD (Baran et al. 2012), or alternative methodological approaches based on neural networks (Montserrat et al. 2020; Oriol Sabat et al. 2022) on our simulated demographic scenarios, could be helpful for researchers interested in applying LAI for particular time periods, sampling schemes and dataset sizes. Although phasing and imputation in ancient genomes using state-of-the-art methods has shown high accuracy (Sousa Da Mota et al. 2023), future follow up work could build on the benchmarking framework established here to evaluate the impact of potential phasing and imputation errors on LAI accuracy. Finally, we note that although our simulations focus on specific values of recombination and mutation rates, these represent adjustable parameters in our reproducible benchmarking pipeline and can thus be extended to other scenarios not covered here.

Our study presents an extensive framework to assess the accuracy of LAI tools, drawing from the comparison between inference and ground truth on simulated data, to provide recommendations with regards to the range of admixture times, sampling strategies, and population sizes which lead to reliable inference. Additionally, we demonstrate the potential of modern simulation methods to carry out extensive reproducible benchmarks applicable to a wide range of population genomic inference tools. With the growing availability of ancient genomic datasets and ongoing advances in statistical inference, our study paves the way for deeper and more accurate insights into the history of past admixture events and their evolutionary consequences.

## Acknowledgments

We thank the members of the Racimo group for their helpful discussions. F.R. and K.B. were supported by a Villum Young Investigator Grant (project no. 00025300). F.R., E.K.I.P. and M.P. were supported by a Novo Nordisk Fonden Data Science Ascending Investigator Award (NNF22OC0076816). E.K.I.-P. was additionally supported by the Royal Society (URF\R1\251675), the European Research Council (ERC) under the European Union’s Horizon Europe programme (grant agreement no. 101222614). F.R. was additionally supported by the European Research Council (ERC) under the European Union’s Horizon Europe and Horizon 2020 programmes (grant agreements No. 101077592 and 951385).

## Author contributions

K.B. and M.P. led the study. K.B., M.P., E.K.I.P., L.A.F.F. and F.R. conceptualized the study. M.P., E.K.I.P., and F.R. supervised the research. F.R. acquired funding for research. K.B. and M.P. undertook formal analysis of the data. K.B. and M.P. drafted the main text. All authors edited the final manuscript.

## Code availability

The code used for the simulations and analyses shown in this study is available at https://github.com/katiabou/LAI_painting.

## Supplementary figures

**Fig. S1:**
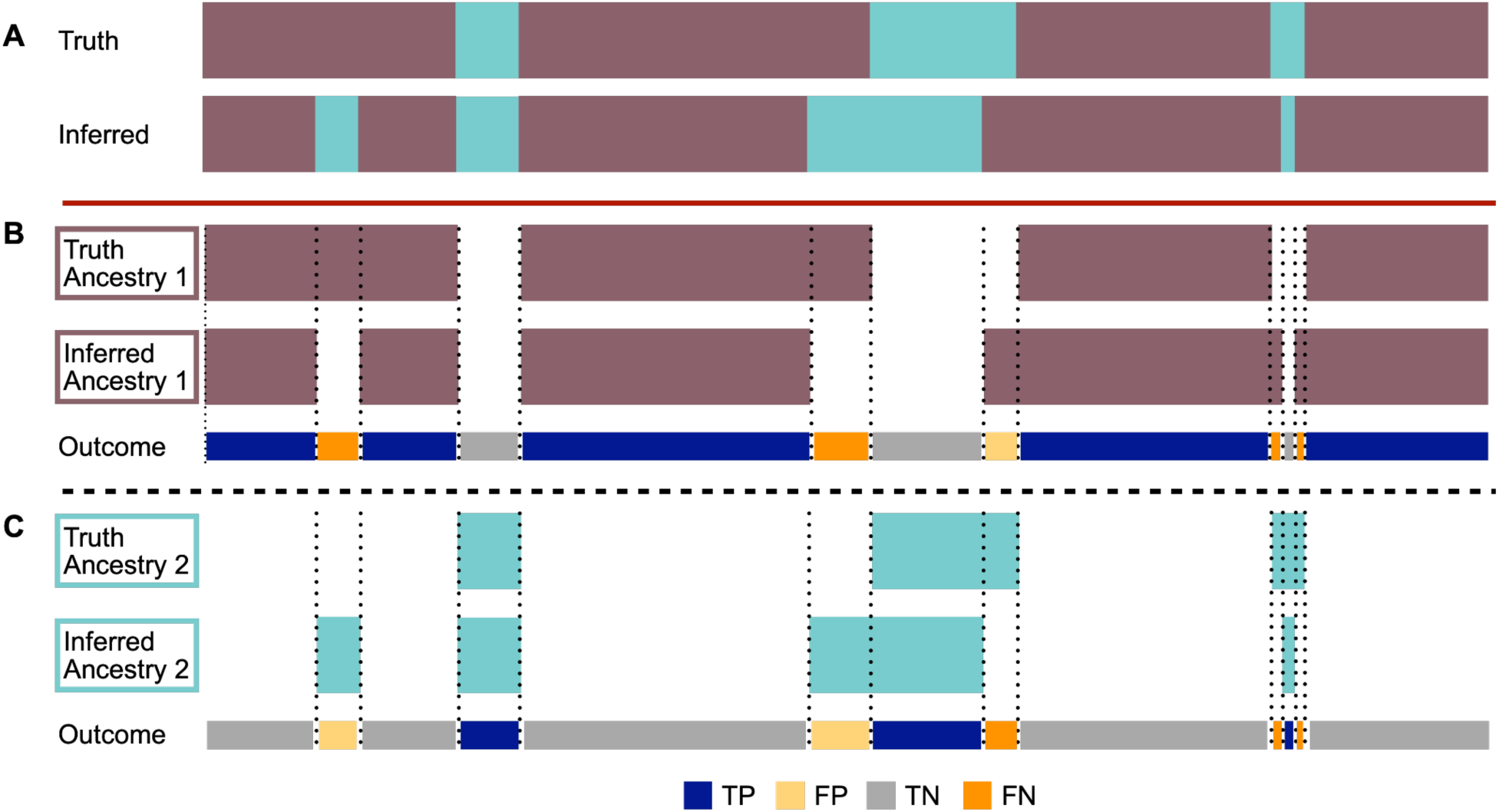
Schematic showcasing the classification of inferred ancestral tracts for one haplotype. A) True ancestral tracts and ancestral tracts inferred by a LAI method. Purple represents ancestry 1 and blue ancestry 2. B) True and inferred ancestral tracts for Ancestry 1 along with the outcome comparison to the known true tracts. C) True and inferred ancestral tracts for Ancestry 2 along with the outcome. Possible outcomes: Dark blue: True Positives (TP), yellow: False Positives (FP), grey: True Negatives (TN), orange: False Negatives (FN). We note the complementary nature of the true tracts (TP and TN) and false tracts (FP and FN) between the two ancestries where, for instance, a TP outcome in one ancestry implies a TN outcome in the other ancestry.

**Fig. S2:**
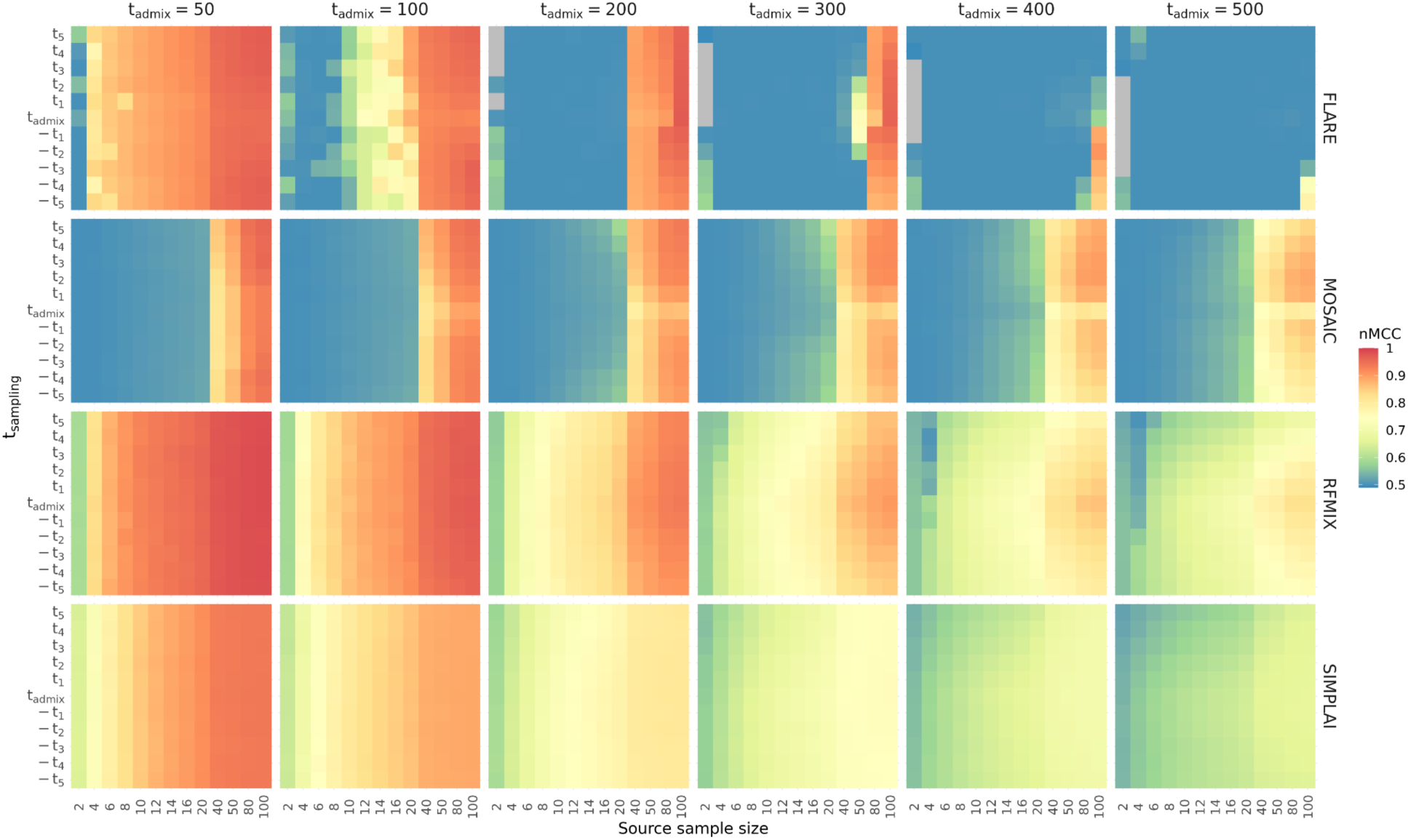
Estimated nMCC metric showcasing the overlap of the inferred and true tracts for the four methods. Due to the complementary nature of all four confusion matrix categories (TP, FP, TN, FN), the nMCC values are the same for the tracts assigned to either of the two ancestries. The y-axis represents the 11 time points at which the source samples were taken from (t_sampling_), relative to the admixture event time (t_admix_). The x-axis is the number of samples taken from each source population. Each column represents the six tested admixture times (t_admix_), using a gene-flow rate of 10%, and each row the results for the four methods. Each square shows the average across all 10 samples from the admixed population and across all 10 replicates. An nMCC value equal to 0.5 (blue) indicates a random prediction and a value closer to 1 (red) represents high overlap. Grey indicates cases where at least one individual from the target population was inferred to have ancestry tracts that did not match any true tracts (TP = 0, FP ≠ 0).

**Fig. S3:**
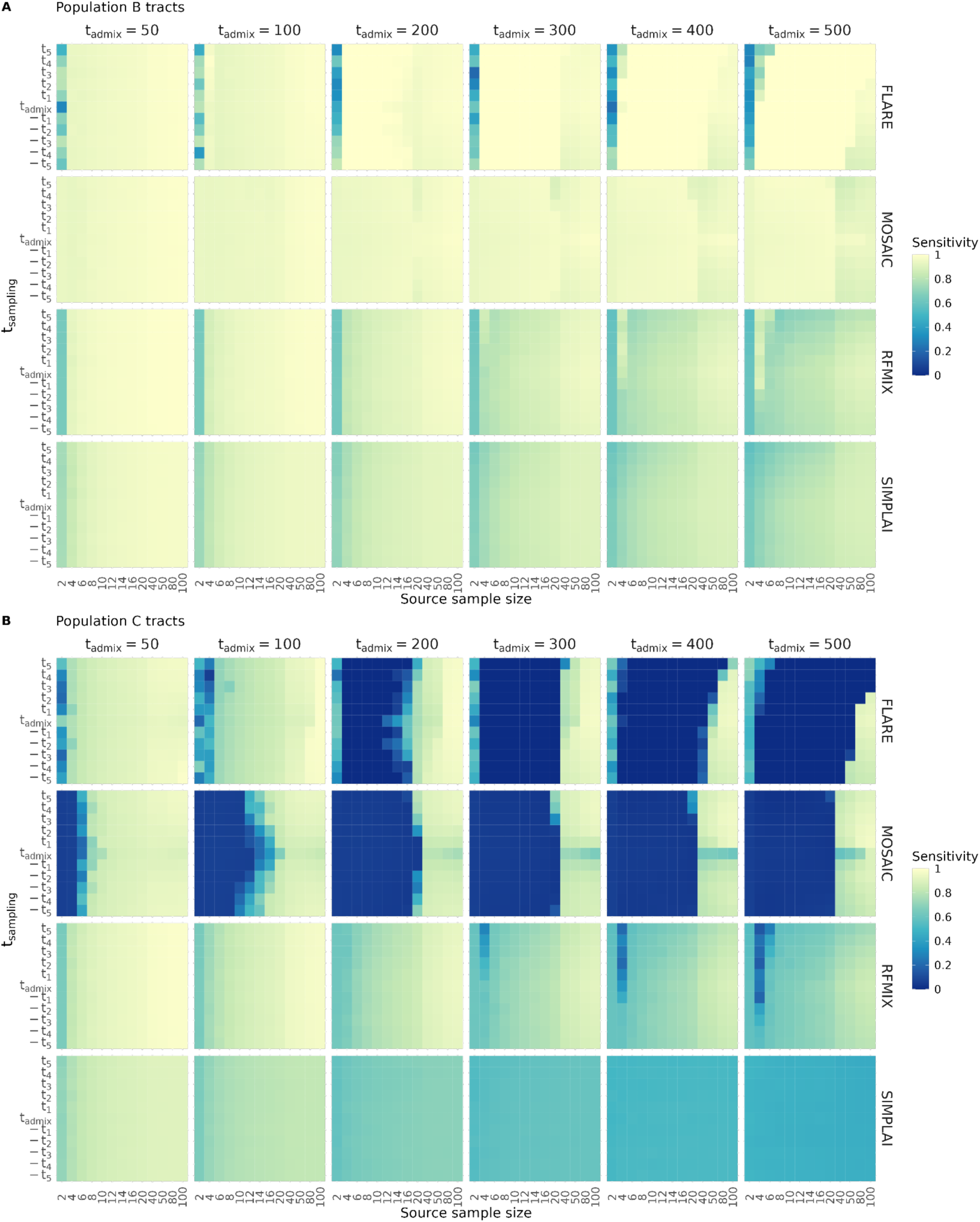
Sensitivity estimates for tracts assigned to A) population B and B) population C across demographic scenarios with a gene-flow rate of 30%. The y-axis represents the 11 time points at which the source samples were taken from (t_sampling_), relative to the admixture event time (t_admix_). The x-axis is the number of samples taken from each source population. Each column represents the five tested admixture times (t_admix_) and each row the results for the four methods. Each square shows the average across all 10 samples from the admixed population and across all 10 replicates. Higher sensitivity estimates represent higher true positive rates.

**Fig. S4:**
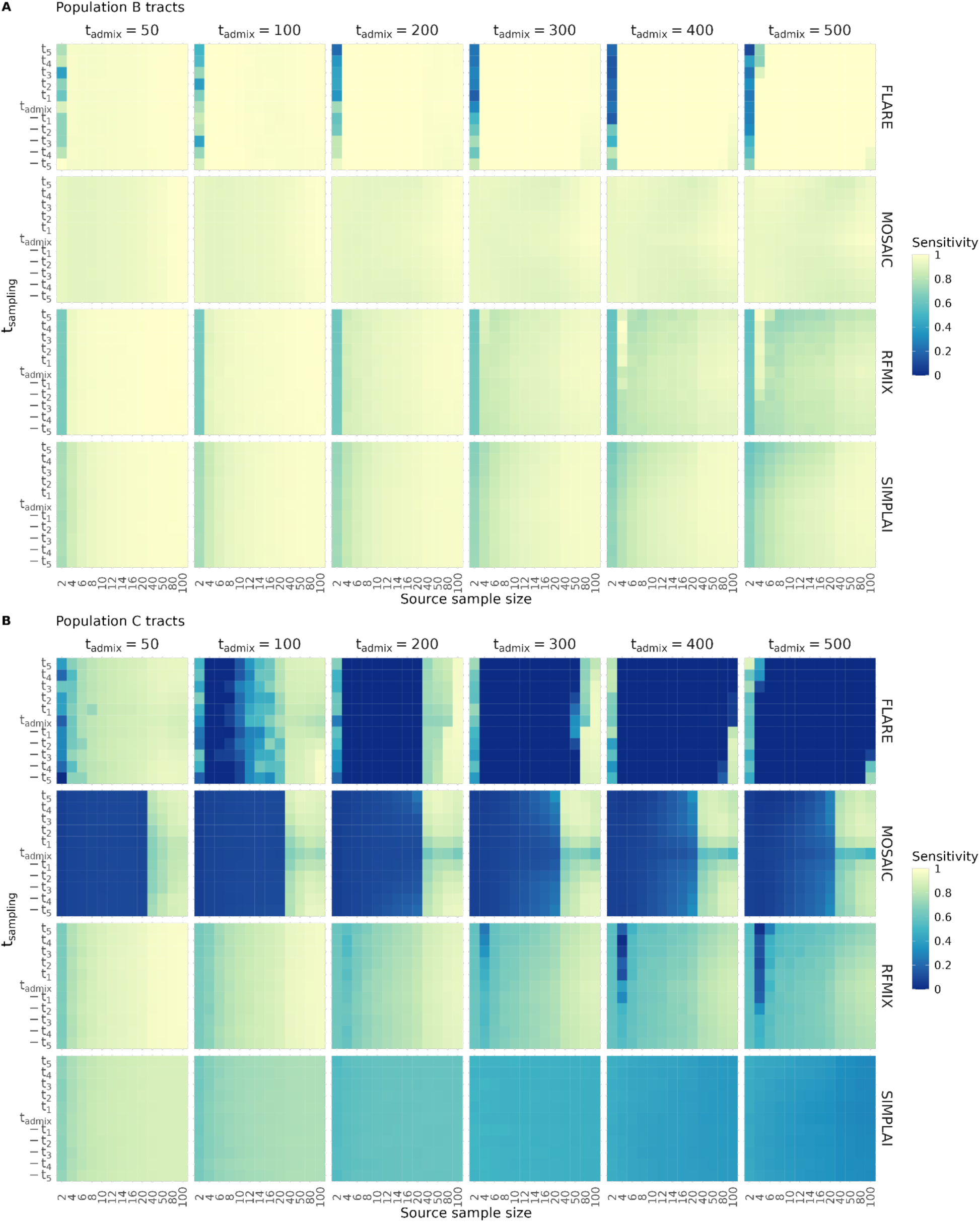
Sensitivity estimates for tracts assigned to A) population B and B) population C across demographic scenarios with a gene-flow rate of 10%. The y-axis represents the 11 time points at which the source samples were taken from (t_sampling_), relative to the admixture event time (t_admix_). The x-axis is the number of samples taken from each source population. Each column represents the five tested admixture times (t_admix_) and each row the results for the four methods. Each square shows the average across all 10 samples from the admixed population and across all 10 replicates. Higher sensitivity estimates represent higher true positive rates.

**Fig. S5:**
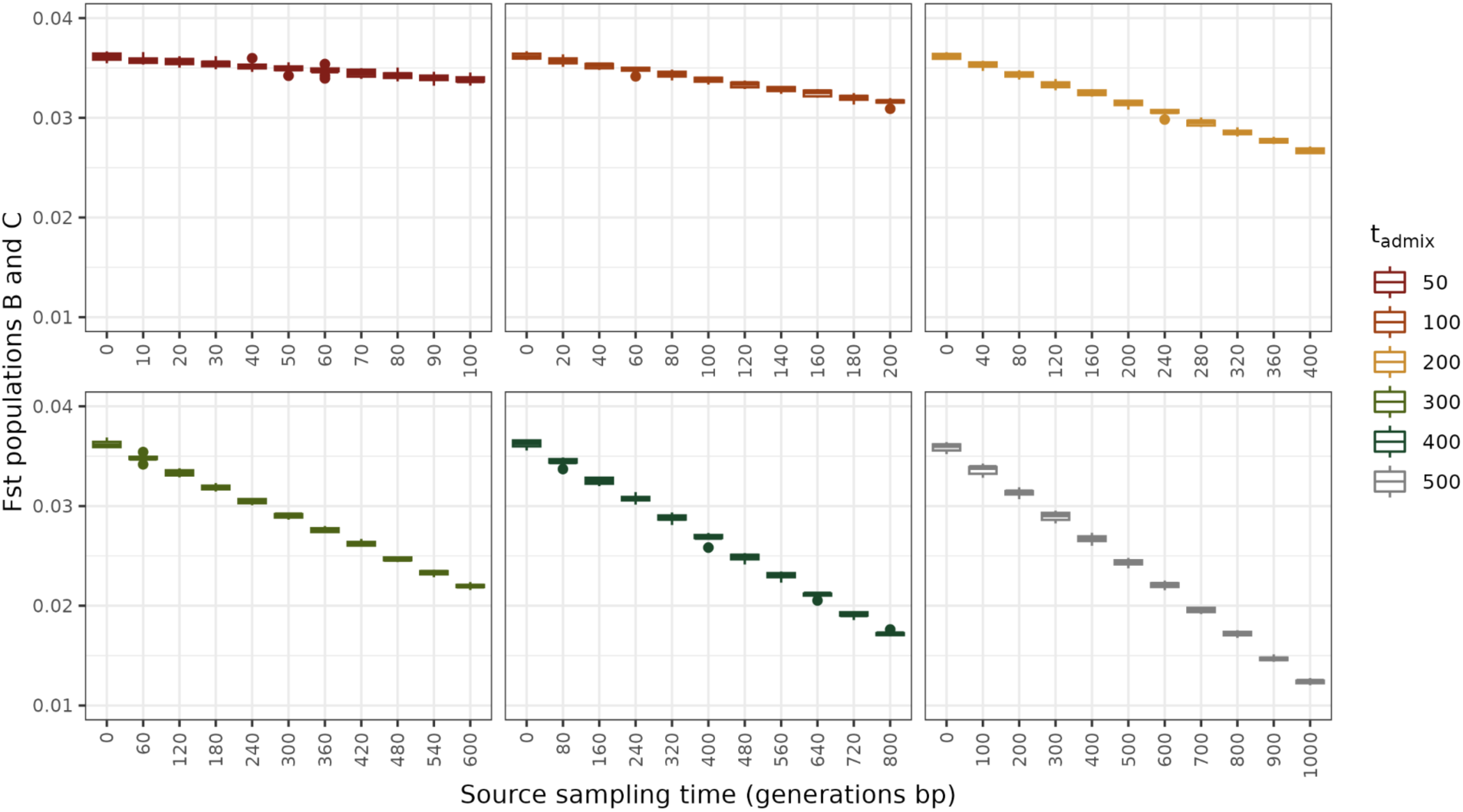
F_ST_ estimates between source populations B and C across admixture times and source sampling time points.

**Fig. S6:**
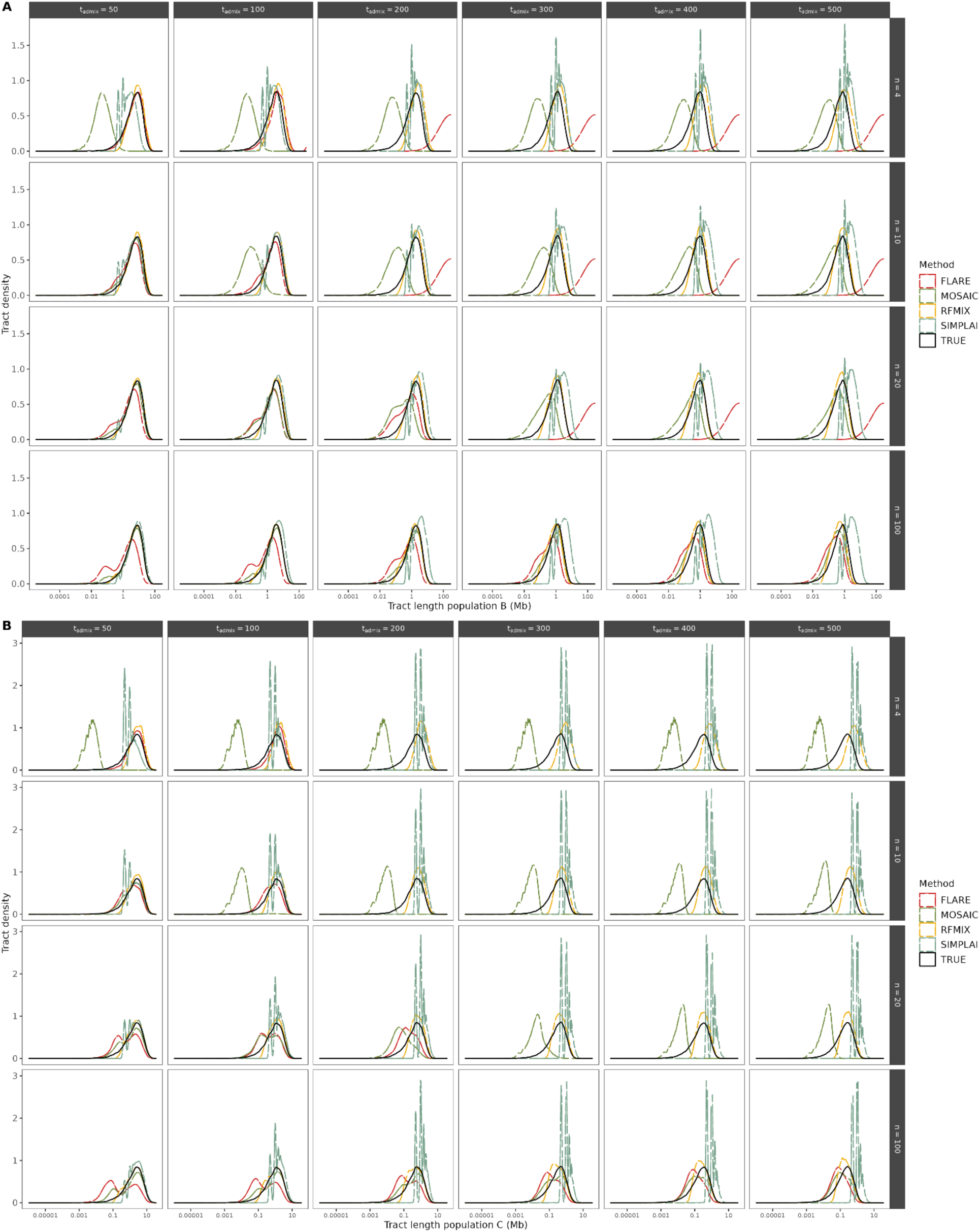
Tract length density plot for true and inferred tracts assigned to A) population B and B) population C across a subset of demographic scenarios with a gene-flow rate of 30%.

**Fig. S7:**
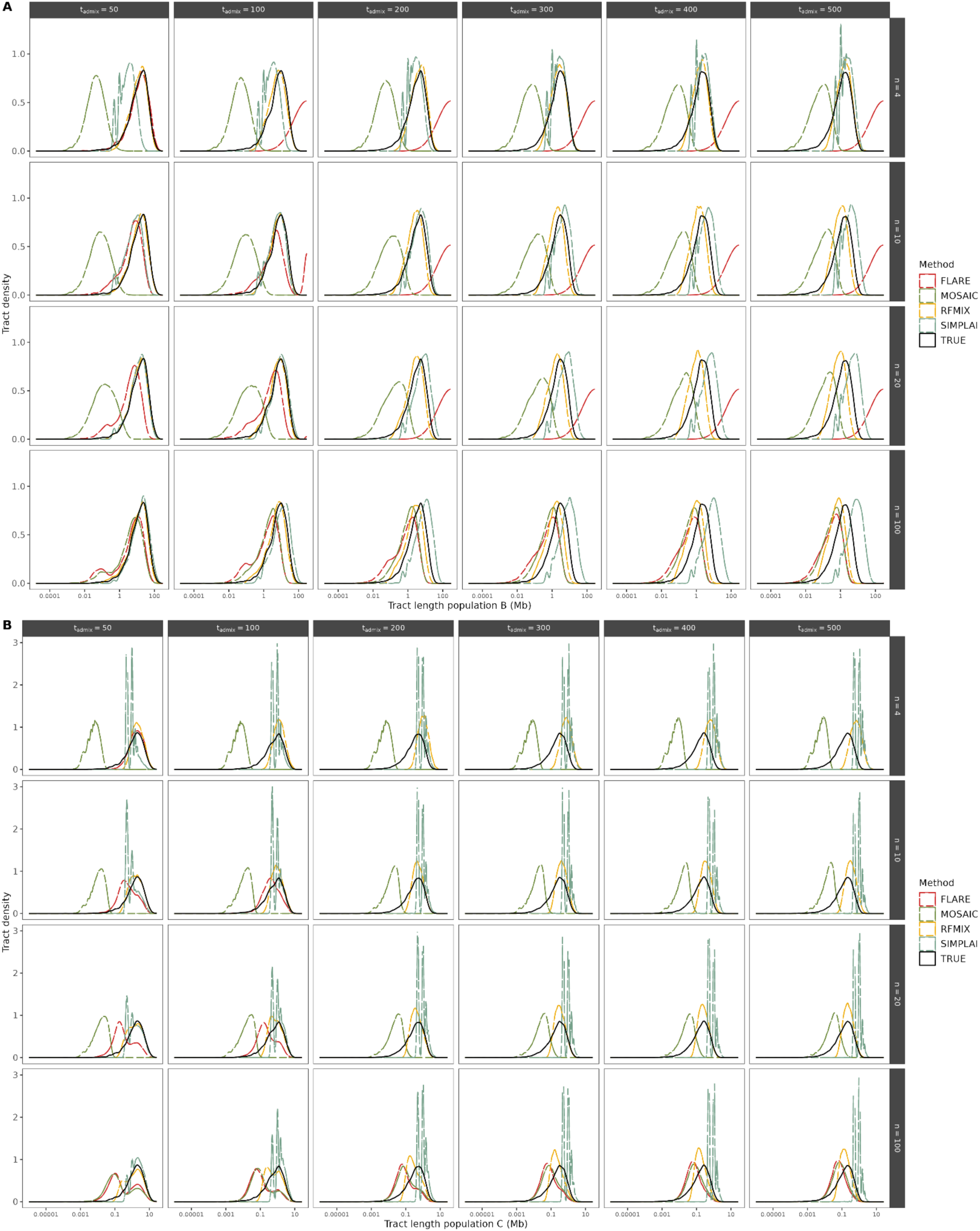
Tract length density plot for true and inferred tracts assigned to A) population B and B) population C across a subset of demographic scenarios with a gene-flow rate of 10%.

**Fig. S8:**
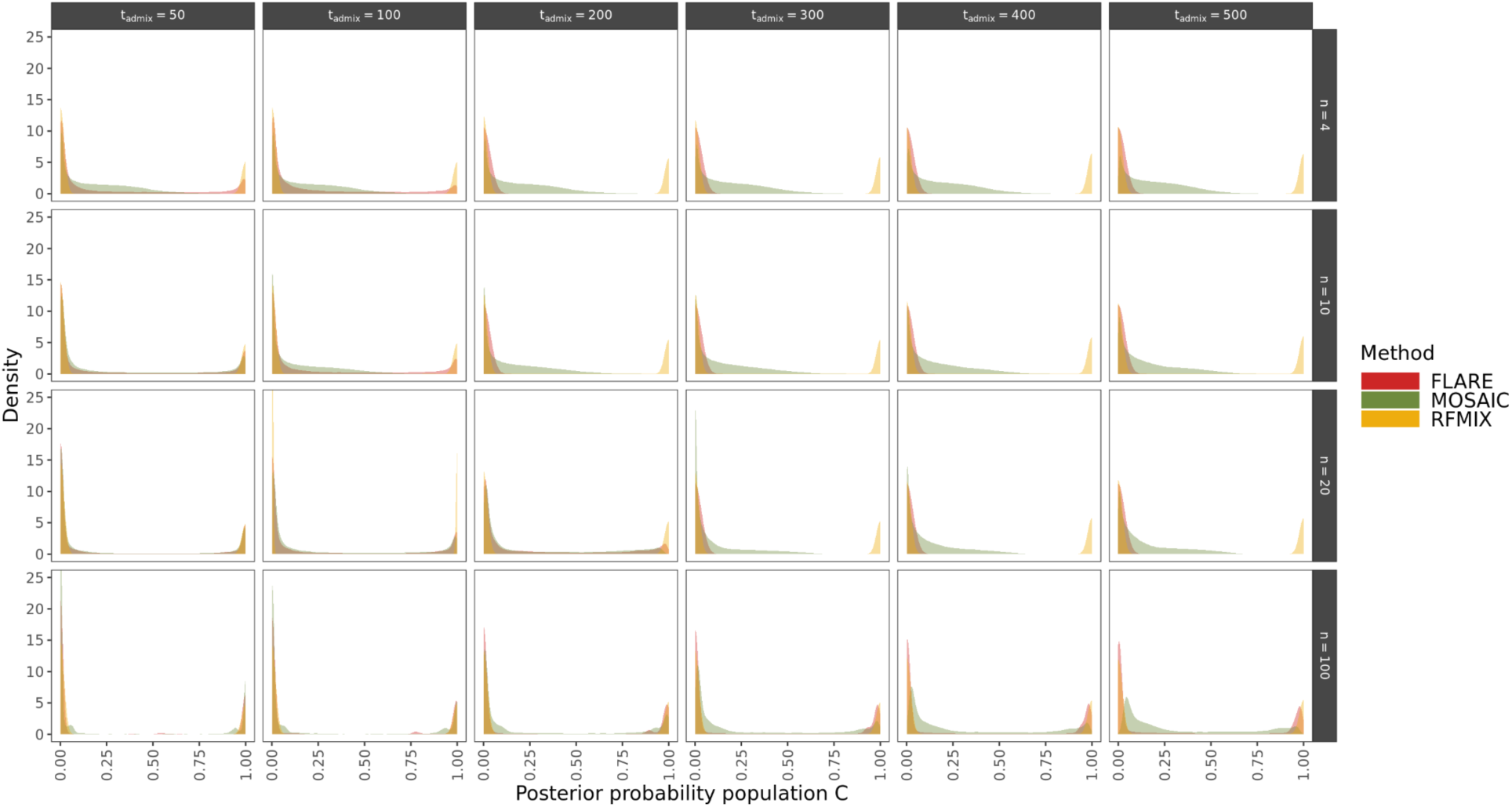
Posterior probability density plot for inferred tracts assigned to population C. Each column represents the five tested admixture times (t_admix_), using a gene-flow rate of 30%, and each row a subset of four source sample sizes (n=4,10,20,100). Results are shown using one source sampling time (t_sampling_=0 generations before present), one seed and a gene-flow rate of 30%.

**Fig. S9:**
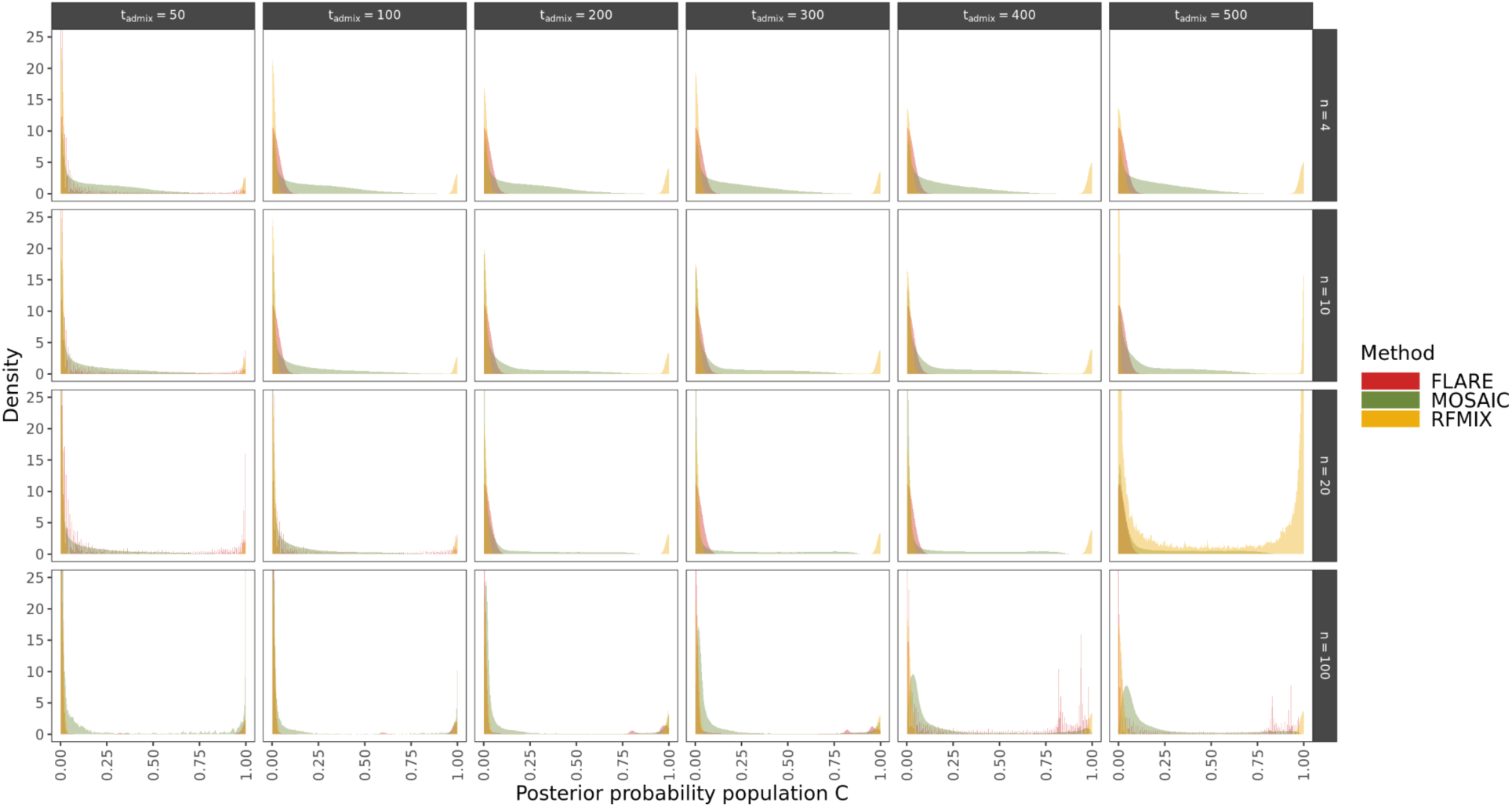
Posterior probability density plot for inferred tracts assigned to population C. Each column represents the five tested admixture times (t_admix_), using a gene-flow rate of 30%, and each row a subset of four source sample sizes (n=4,10,20,100). Results are shown using one source sampling time (t_sampling_=0 generations before present), one seed and a gene-flow rate of 10%.

**Fig. S10:**
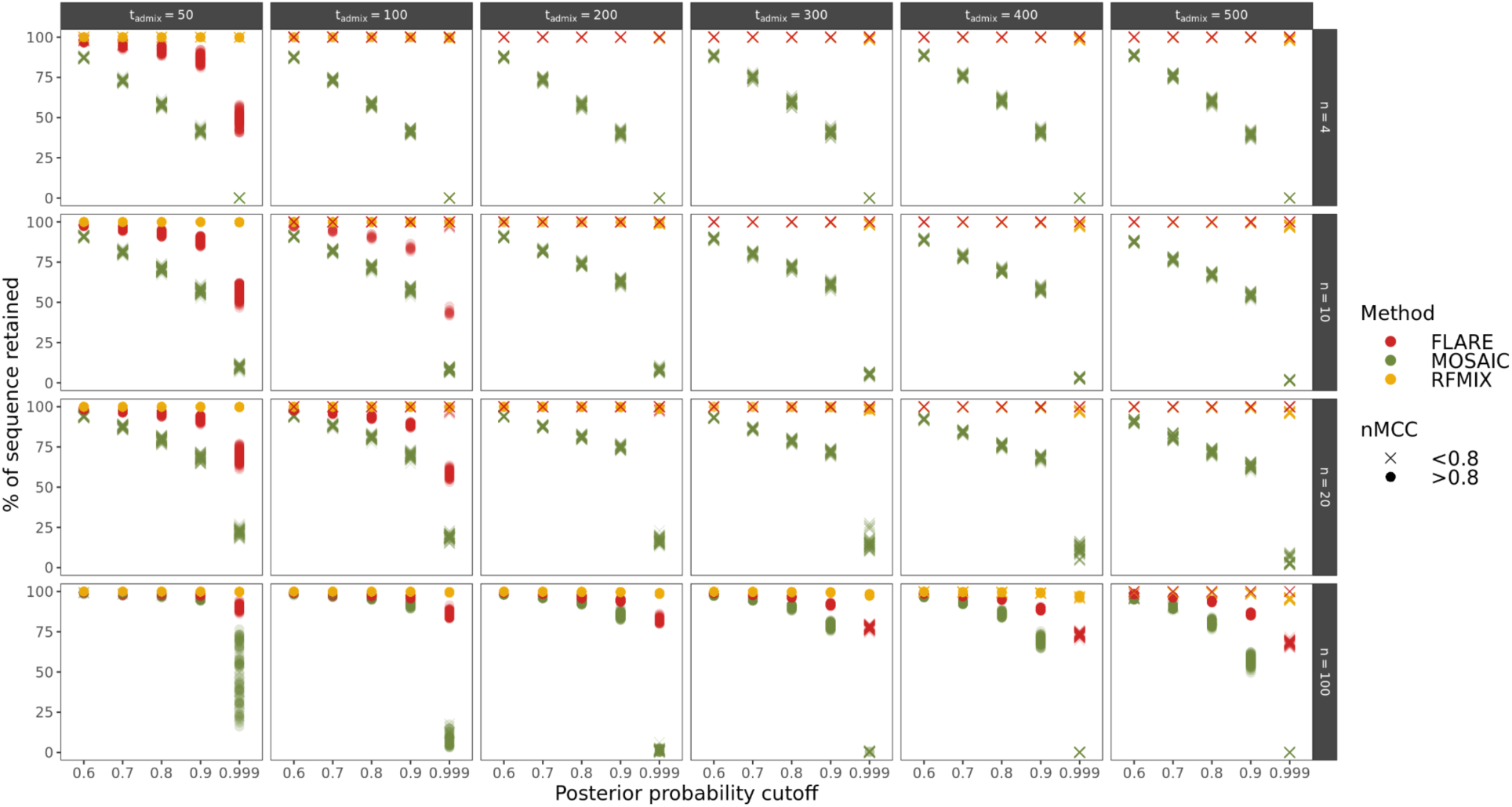
Percentage of sequence left after applying five different probability cutoffs. Each column represents the five tested admixture times (t_admix_), using a gene-flow rate of 10%, and each row a subset of four source sample sizes (n=4,10,20,100). Results are shown for one source sampling time (t_sampling_=0 generations before present). The colour represents the method used and shape depicts nMCC values either above or below 0.8. Each point shows the estimated value for each of the 10 target samples from each of the 10 replicates. We note that simpLAI is missing in this comparison because it reports ancestry tracts for the entire genome without any posterior probability output.

**Fig. S11:**
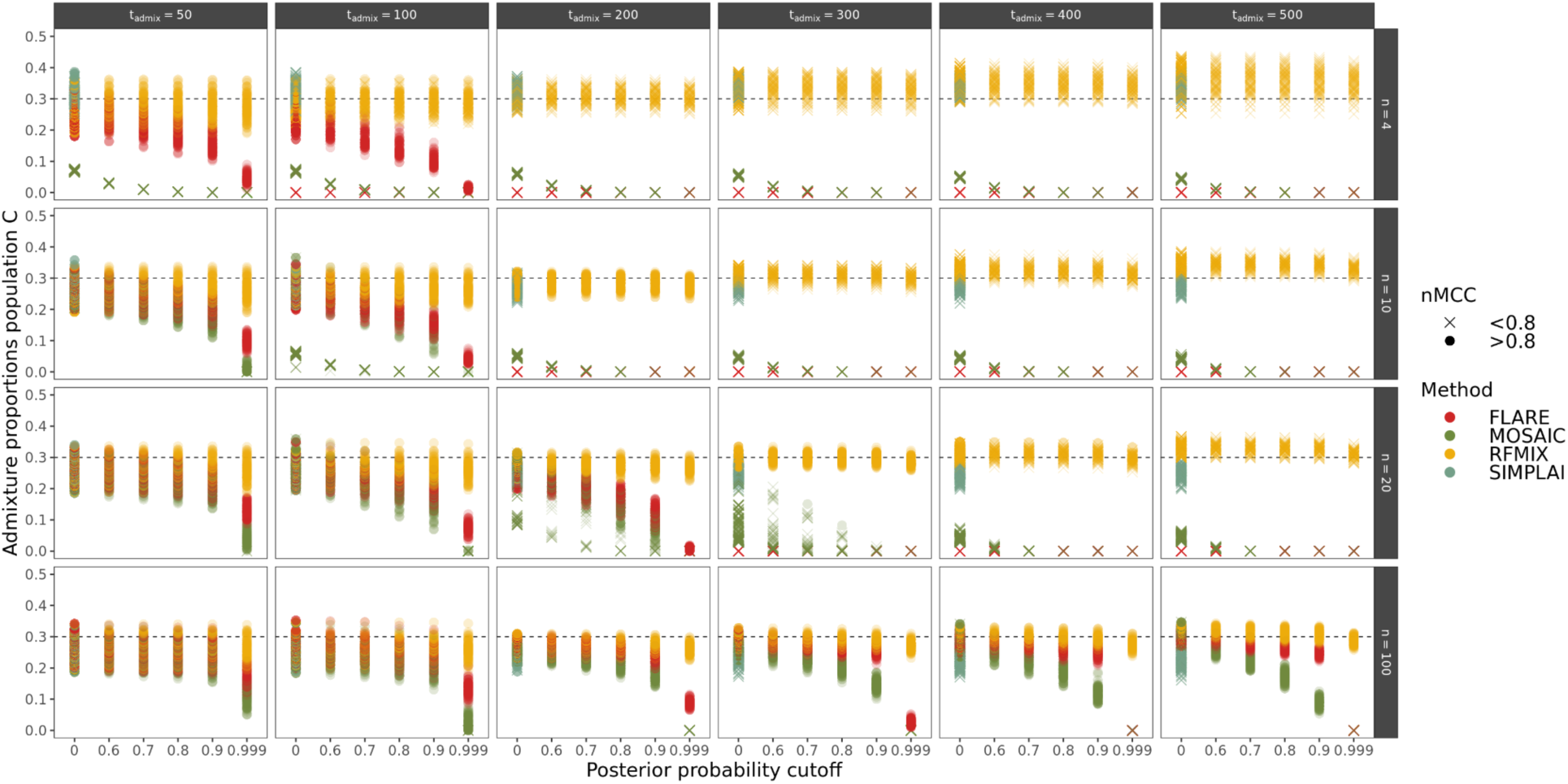
Estimated admixture proportions for population C based on the sequence left after applying five different probability cutoffs. Each column represents the five tested admixture times (t_admix_), using a gene-flow rate of 30%, and each row a subset of four source sample sizes (n=4,10,20,100). Results are shown for one source sampling time (t_sampling_=0 generations before present). The colour represents the method used and shape depicts nMCC values either above or below 0.8. Each point shows the estimated value for each of the 10 target samples from each of the 10 replicates. We note that simpLAI is only present in the case with no posterior probability cutoff as it reports ancestry tracts for the entire genome without any posterior probability output.

**Fig. S12:**
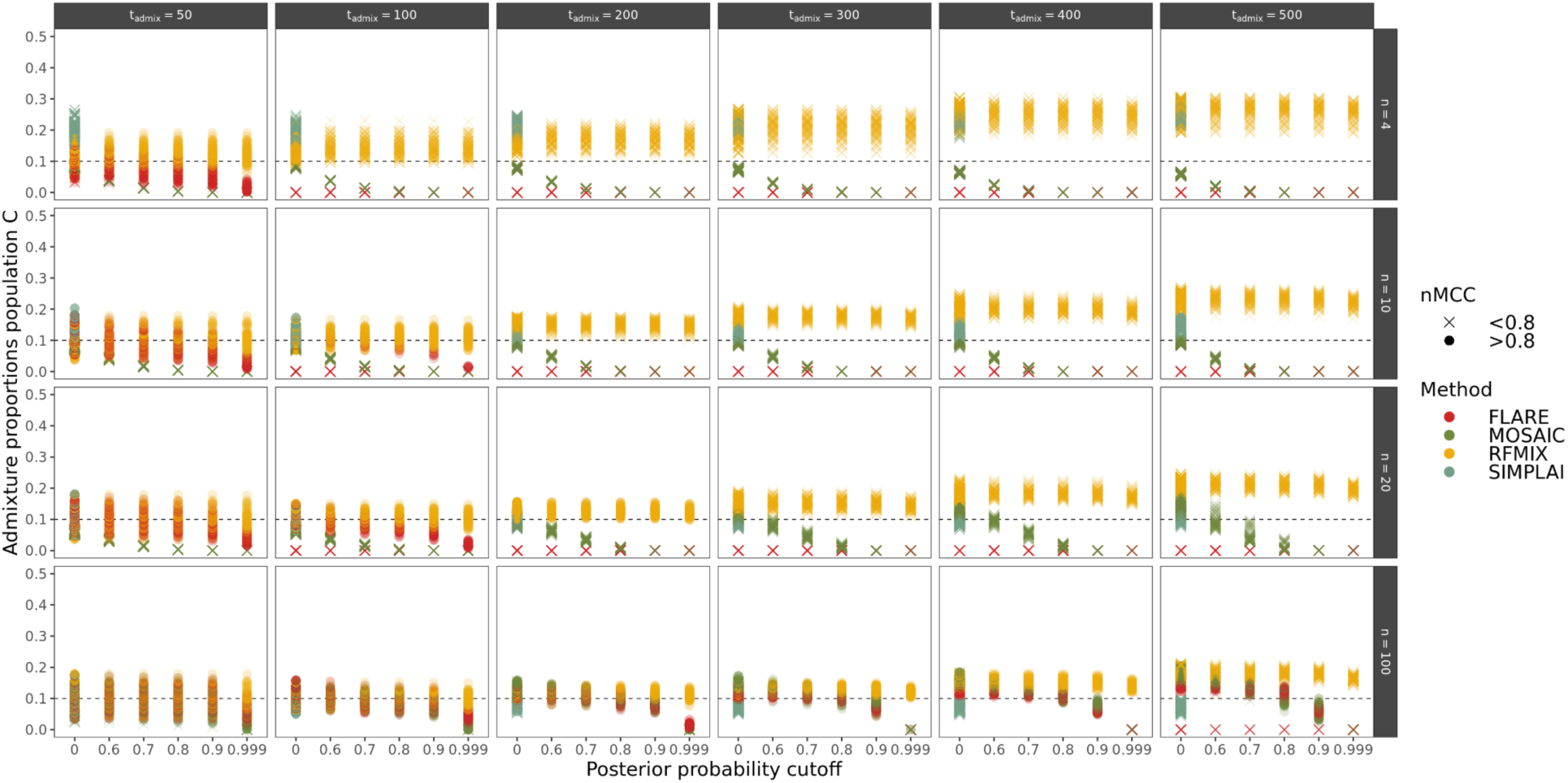
Estimated admixture proportions for population C based on the sequence left after applying five different probability cutoffs. Each column represents the five tested admixture times (t_admix_), using a gene-flow rate of 10%, and each row a subset of four source sample sizes (n=4,10,20,100). Results are shown for one source sampling time (t_sampling_=0 generations before present). The colour represents the method used and shape depicts nMCC values either above or below 0.8. Each point shows the estimated value for each of the 10 target samples from each of the 10 replicates. We note that simpLAI is only present in the case with no posterior probability cutoff as it reports ancestry tracts for the entire genome without any posterior probability output.

**Fig. S13:**
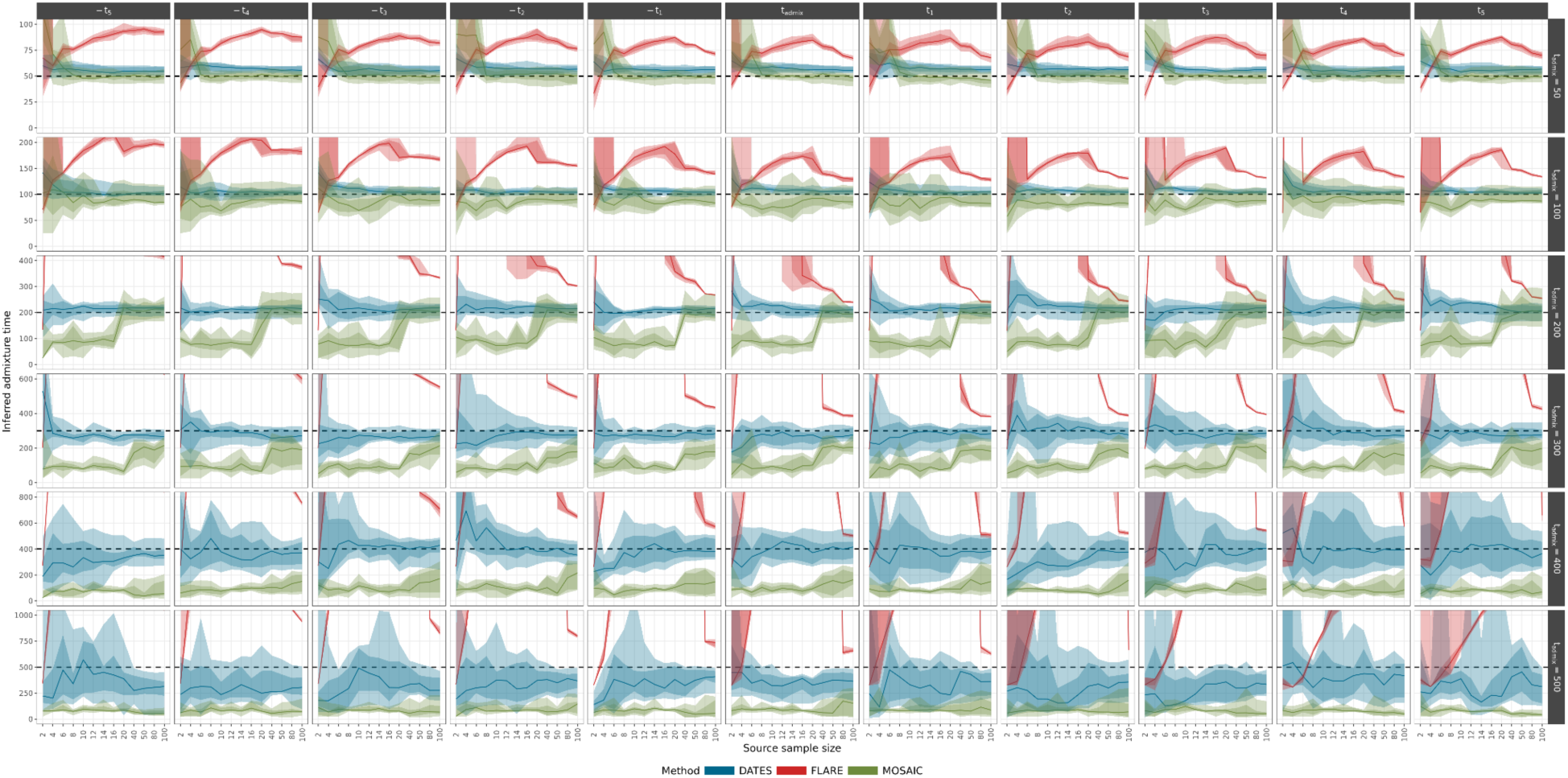
Inferred admixture times (y axis) under each combination of demographic parameters from DATES (blue), FLARE (red) and MOSAIC (green). The x-axis shows the 13 sample sizes taken from each of the two ancestries. Each column represents one of the 11 time points at which the source samples were taken from (t_sampling_) and each row one of the six tested admixture times (t_admix_). All results are shown for a gene-flow rate of 30%. The dotted black line indicates the true admixture time. The shaded regions in each ribbon show the 50% and 80% quantiles and the line represents the median. Estimates which fall outside of the range [0, 2×t_admix_] are not shown.

**Fig. S14:**
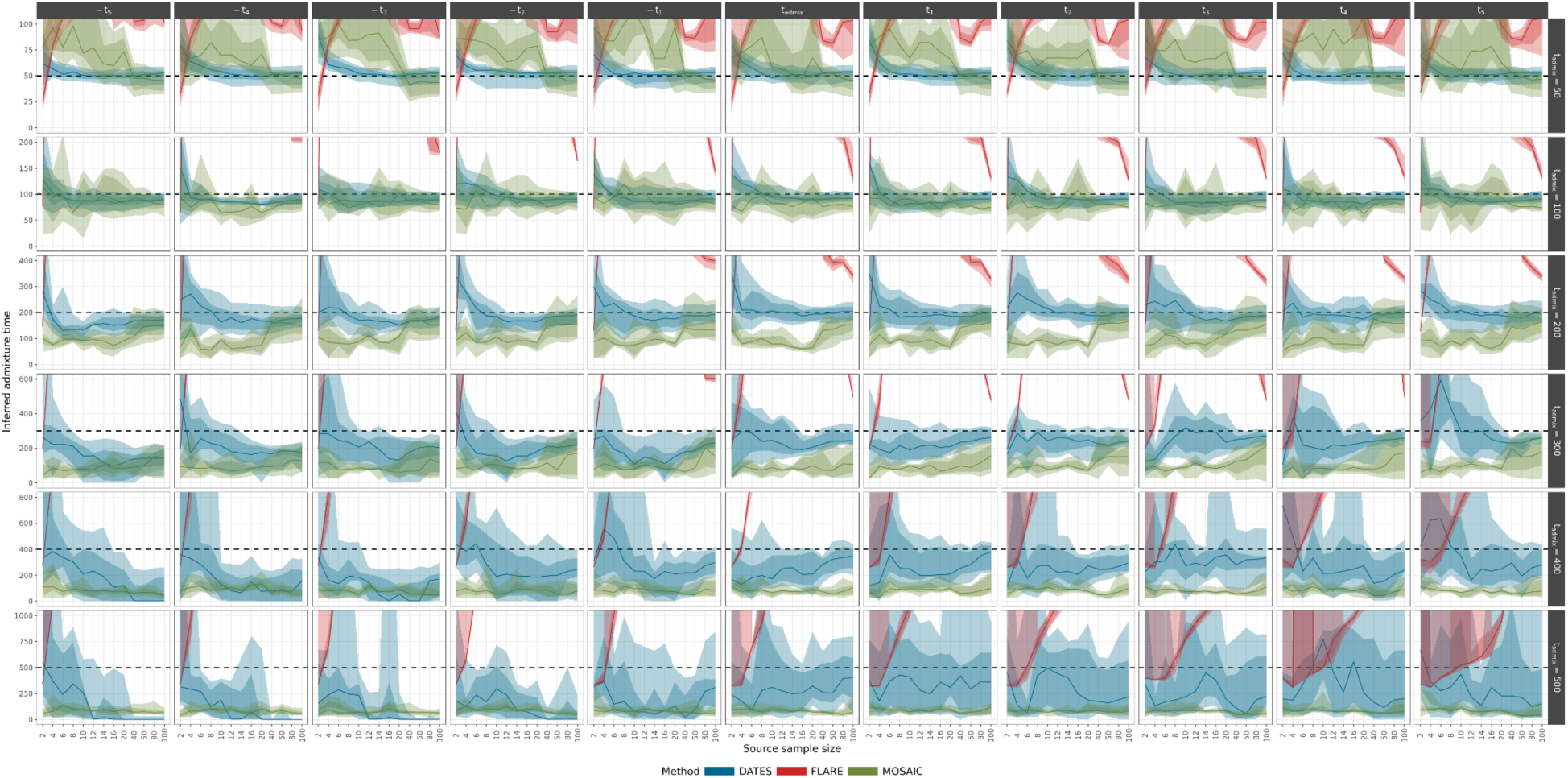
Inferred admixture times (y axis) under each combination of demographic parameters from DATES (blue), FLARE (red) and MOSAIC (green). The x-axis shows the 13 sample sizes taken from each of the two ancestries. Each column represents one of the 11 time points at which the source samples were taken from (t_sampling_) and each row one of the six tested admixture times (t_admix_). All results are shown for a gene-flow rate of 10%. The dotted black line indicates the true admixture time. The shaded regions in each ribbon show the 50% and 80% quantiles and the line represents the median. Estimates which fall outside of the range [0, 2×t_admix_] are not shown.

**Fig. S15:**
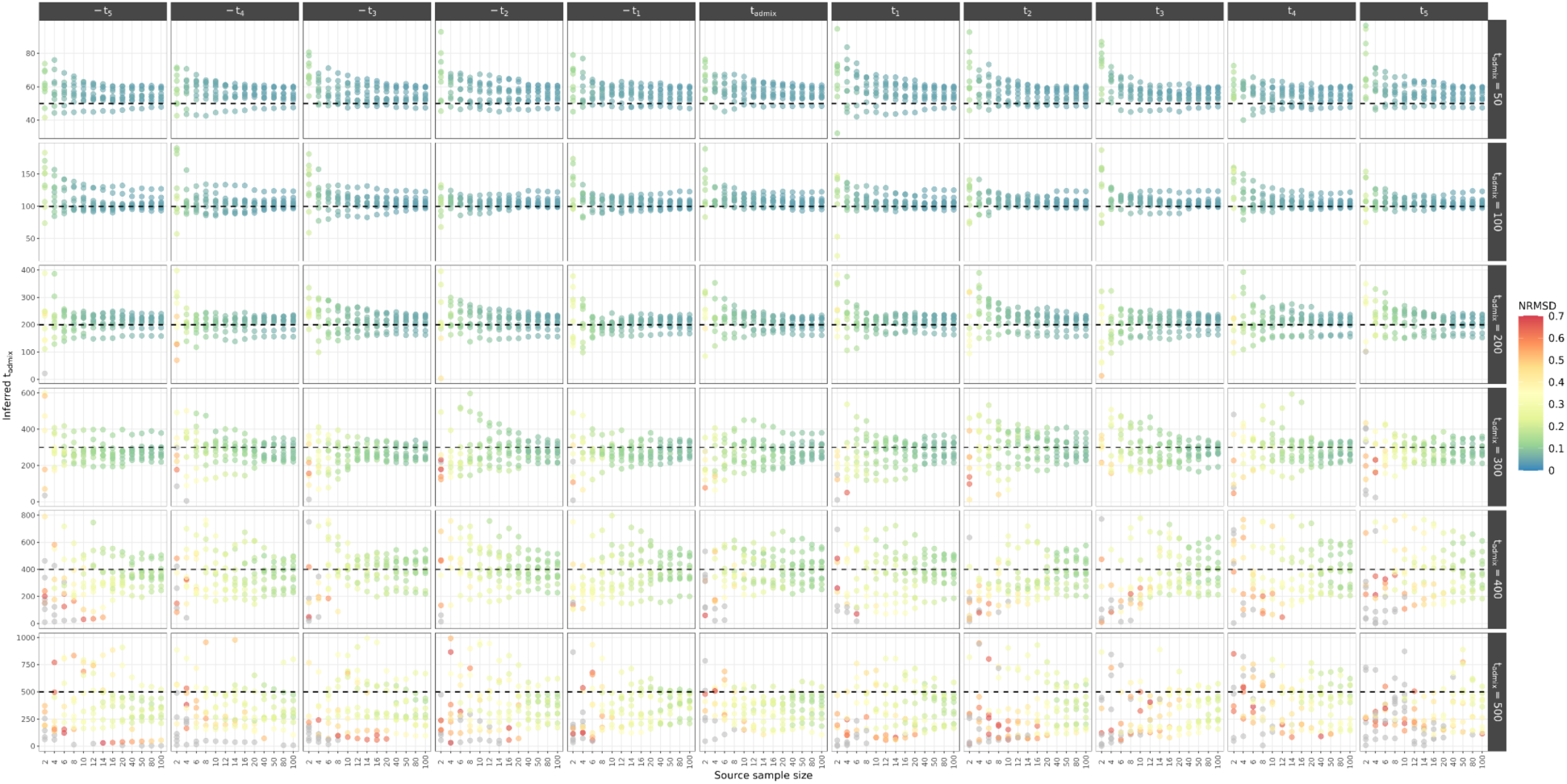
Inferred admixture times (y axis) under each combination of demographic parameters from DATES, coloured by the NRMSD values, with grey indicating estimates above the recommended upper limit of 0.7. The x-axis shows the 13 sample sizes taken from each of the two ancestries. Each column represents one of the 11 time points at which the source samples were taken from (t_sampling_) and each row one of the six tested admixture times (t_admix_). All results are shown for a gene-flow rate of 30%. The dotted black line indicates the true admixture time. Each point shows the estimated value from each of the 10 replicates. Inferred admixture times which were ≥ 2×t_admix_ were removed from the plot to improve clarity.

**Fig. S16:**
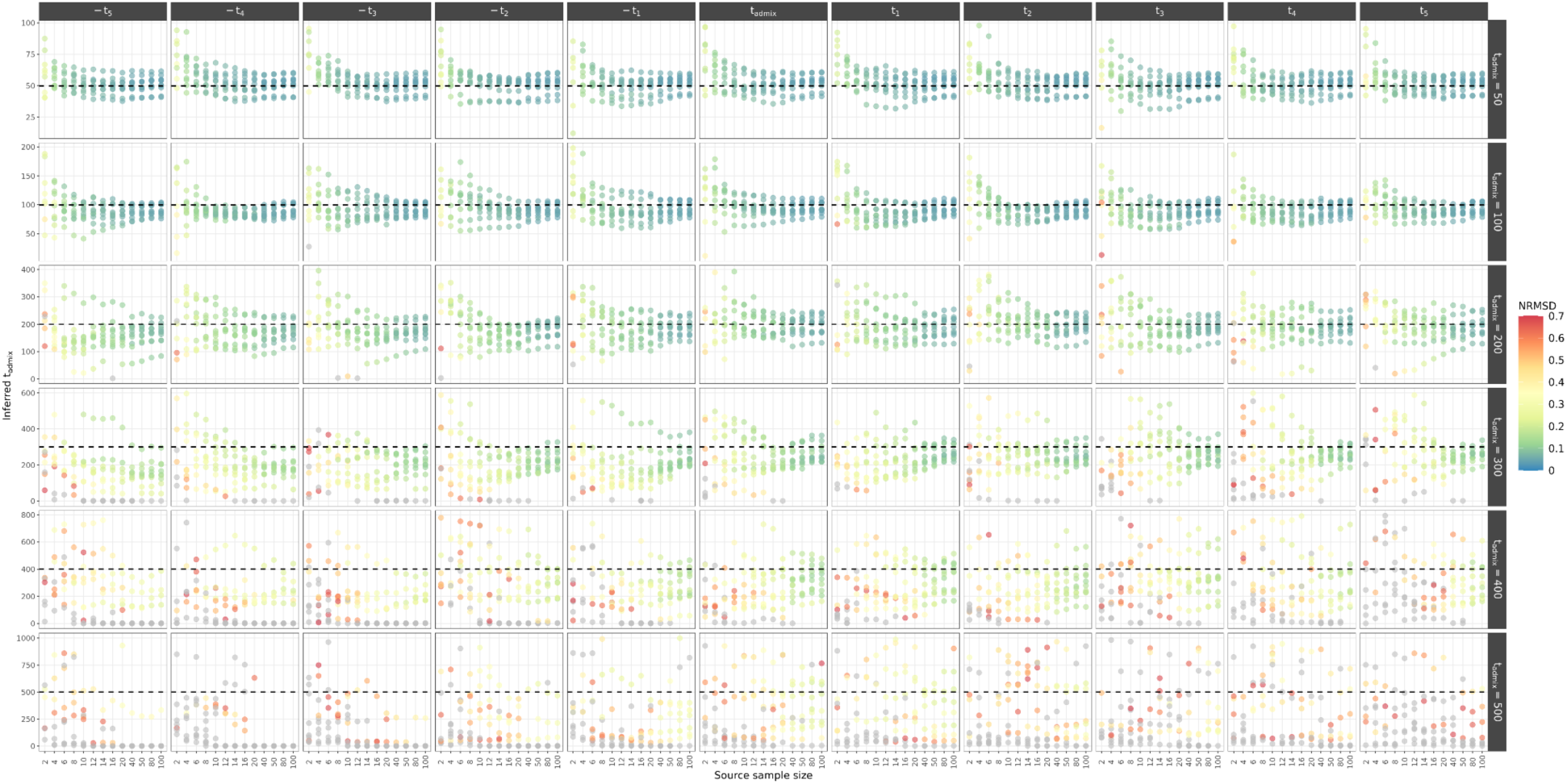
Inferred admixture times (y axis) under each combination of demographic parameters from DATES, coloured by the NRMSD values, with grey indicating estimates above the recommended upper limit of 0.7. The x-axis shows the 13 sample sizes taken from each of the two ancestries. Each column represents one of the 11 time points at which the source samples were taken from (t_sampling_) and each row one of the six tested admixture times (t_admix_). All results are shown for a gene-flow rate of 10%. The dotted black line indicates the true admixture time. Each point shows the estimated value from each of the 10 replicates. Inferred admixture times which were ≥ 2×t_admix_ were removed from the plot to improve clarity.

**Fig. S17:**
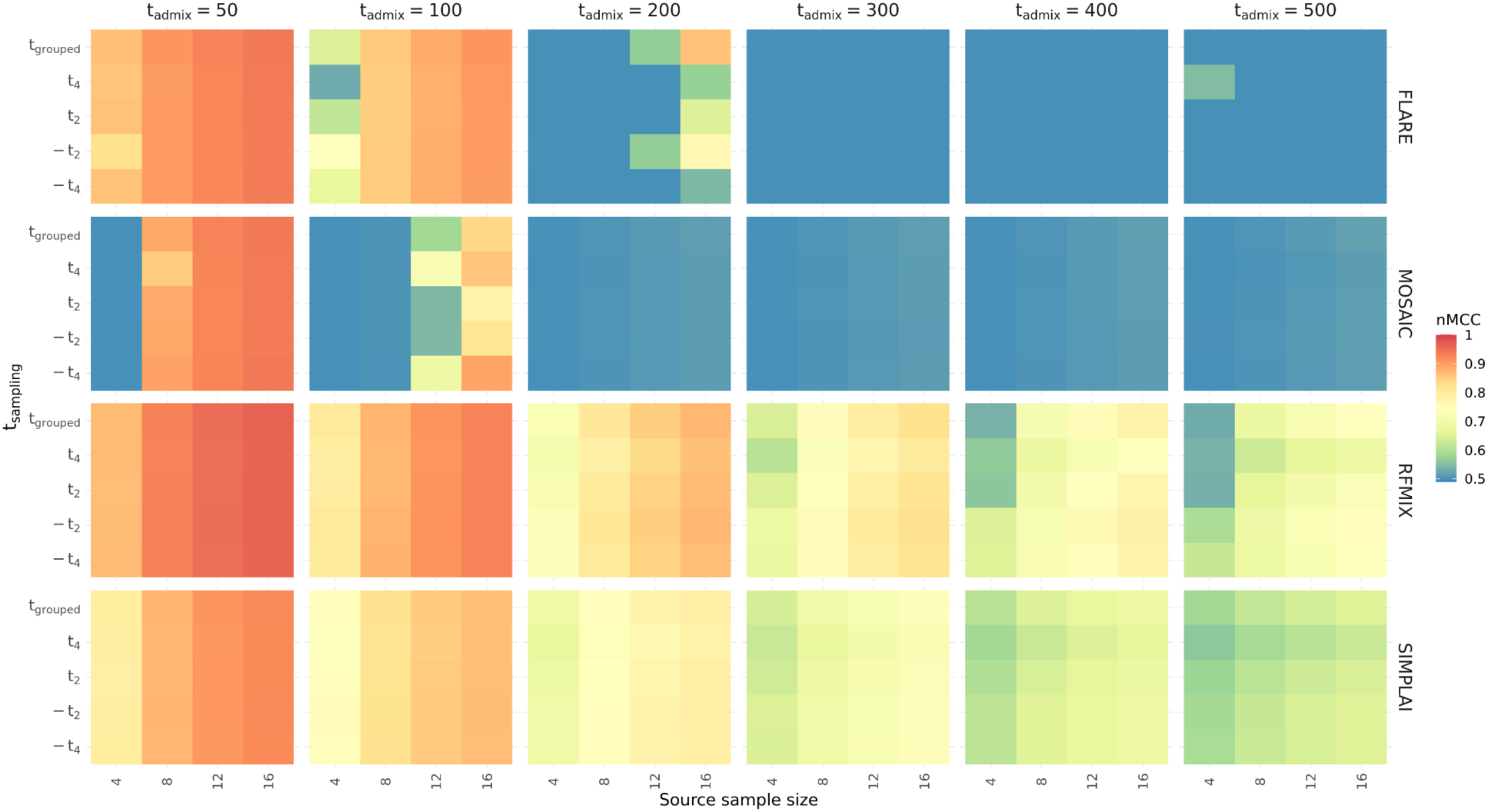
Estimated nMCC metric showcasing the overlap of the inferred and true tracts for the four methods, using a merged time series dataset of sources sampled from four time points (t_sampling_ : t_-4_=t_admix_×0.2, t_-2_=t_admix_×0.6, t_2_=t_admix_×1.4, t_4_=t_admix_×1.8). Due to the complementary nature of all four confusion matrix categories (TP, FP, TN, FN), the nMCC values are the same for the tracts assigned to either of the two ancestries. The y-axis represents the four time points at which the source samples were taken from, with the fifth time point representing the time-series run (t_grouped_). The x-axis shows the four sample sizes taken from each source population. Each column represents the five tested admixture times (t_admix_), using a gene-flow rate of 30%, and each row the results for the four methods. Each square shows the average across all 10 samples from the admixed population and across all 10 replicates. An nMCC value equal to 0.5 (blue) indicates a random prediction and a value closer to 1 (red) represents high overlap.

**Fig. S18:**
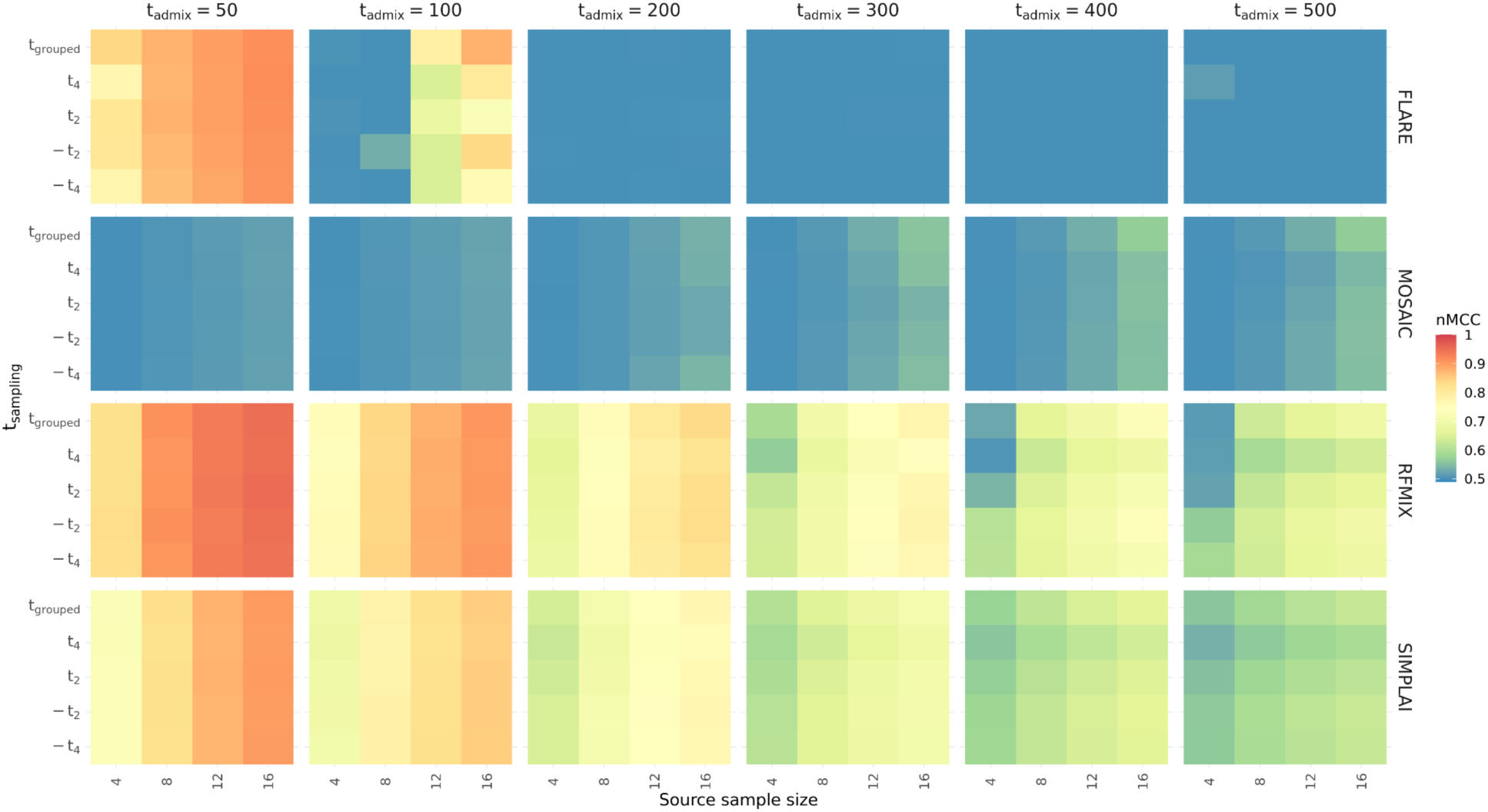
Estimated nMCC metric showcasing the overlap of the inferred and true tracts for the four methods, using a merged time series dataset of sources sampled from four time points (t_sampling_ : t_-4_=t_admix_×0.2, t_-2_=t_admix_×0.6, t_2_=t_admix_×1.4, t_4_=t_admix_×1.8). Due to the complementary nature of all four confusion matrix categories (TP, FP, TN, FN), the nMCC values are the same for the tracts assigned to either of the two ancestries. The y-axis represents the four time points at which the source samples were taken from, with the fifth time point representing the time-series run (t_grouped_). The x-axis shows the four sample sizes taken from each source population. Each column represents the five tested admixture times (t_admix_), using a gene-flow rate of 10%, and each row the results for the four methods. Each square shows the average across all 10 samples from the admixed population and across all 10 replicates. An nMCC value equal to 0.5 (blue) indicates a random prediction and a value closer to 1 (red) represents high overlap.

**Fig. S19:**
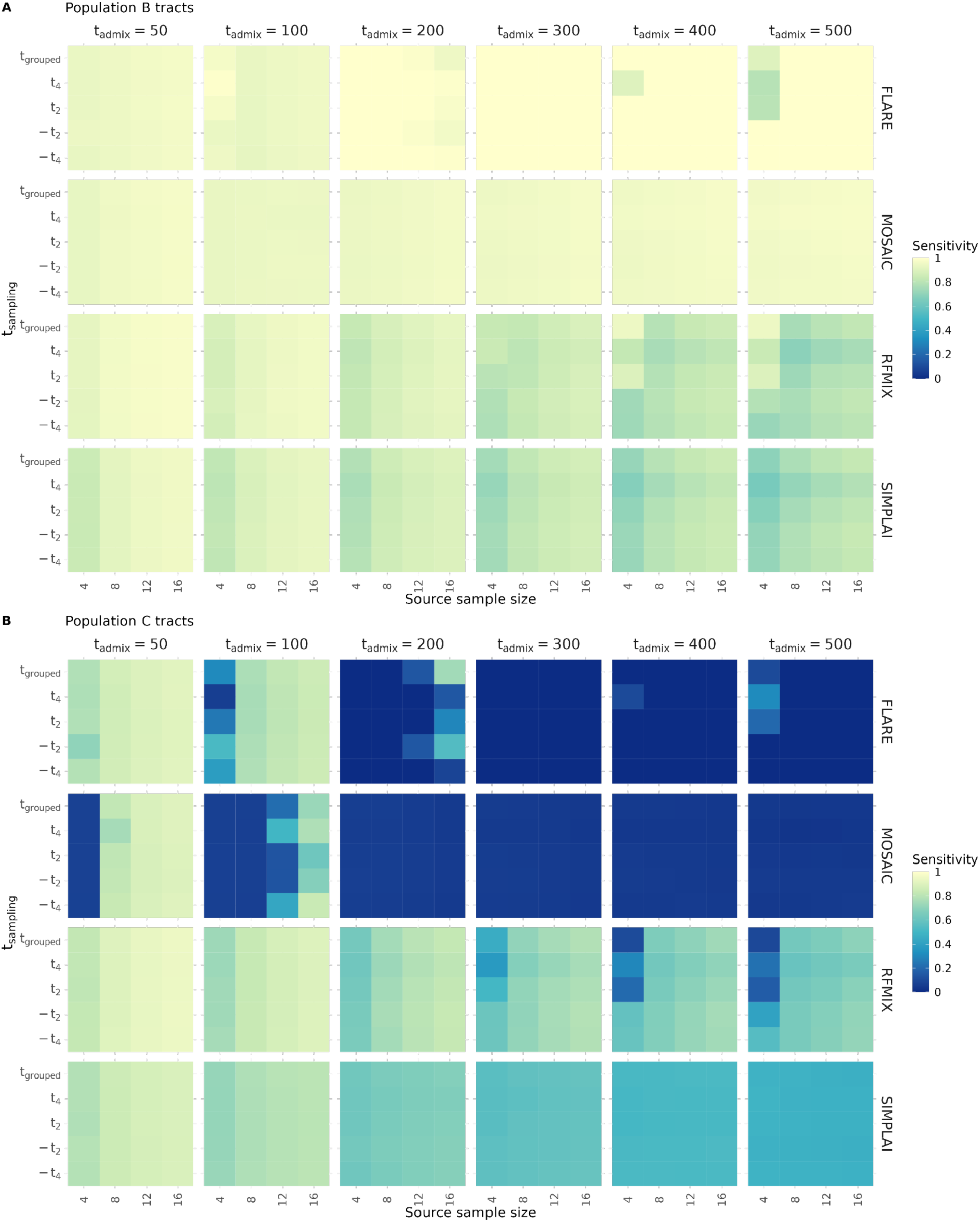
Sensitivity estimates for tracts assigned to A) population B and B) population C, using a merged time series dataset of sources sampled from four time points (t_sampling_ : t_-4_=t_admix_×0.2, t_-2_=t_admix_×0.6, t_2_=t_admix_×1.4, t_4_=t_admix_×1.8). The y-axis represents the four time points at which the source samples were taken from, with the fifth time point representing the time-series run (t_grouped_). The x-axis shows the four sample sizes taken from each source population. Each column represents the five tested admixture times (t_admix_), using a gene-flow rate of 30%, and each row the results for the four methods. Each square shows the average across all 10 samples from the admixed population and across all 10 replicates. Higher sensitivity estimates represent higher true positive rates.

**Fig. S20:**
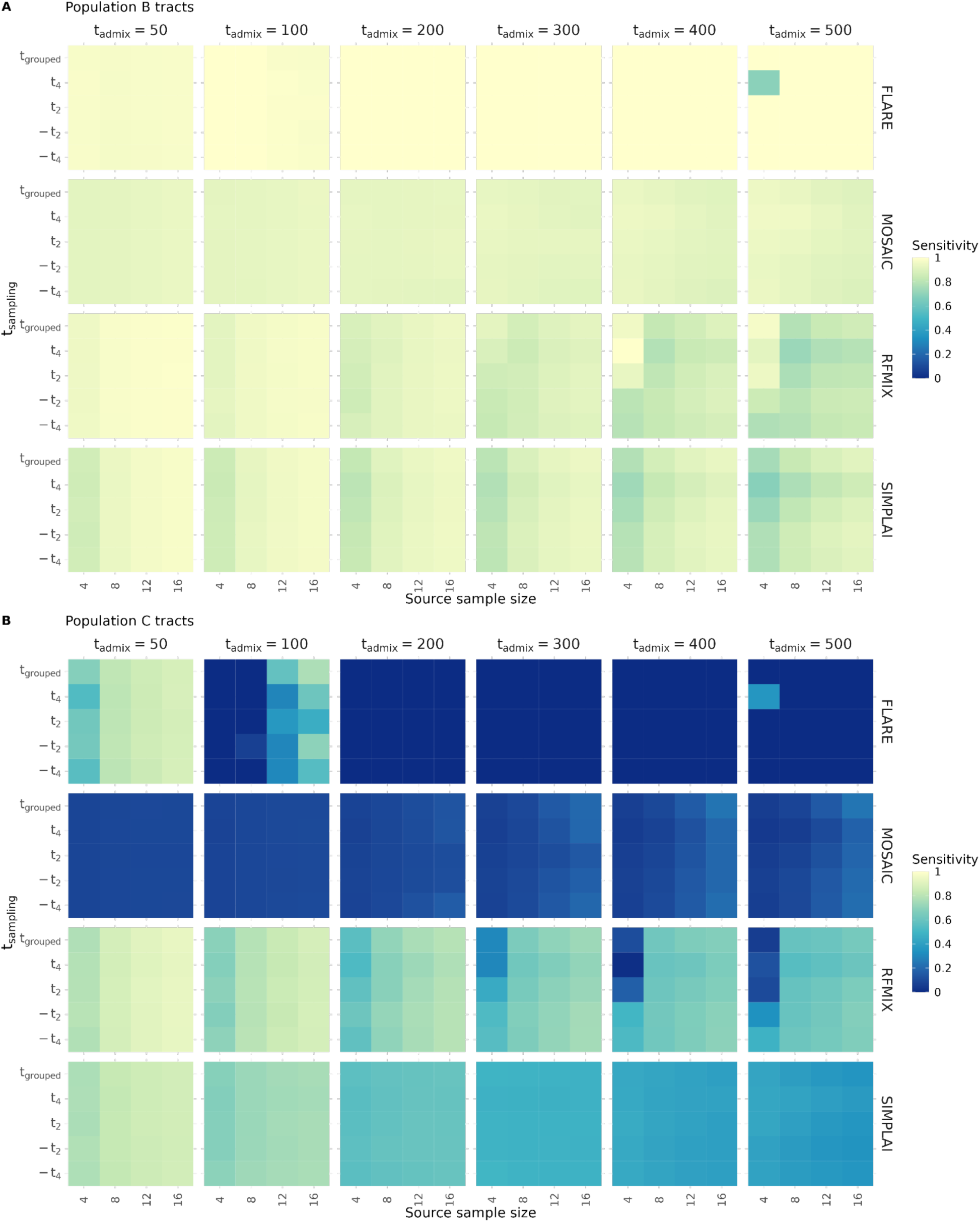
Sensitivity estimates for tracts assigned to A) population B and B) population C, using a merged time series dataset of sources sampled from four time points (t_sampling_ : t_-4_=t_admix_×0.2, t_-2_=t_admix_×0.6, t_2_=t_admix_×1.4, t_4_=t_admix_×1.8). The y-axis represents the four time points at which the source samples were taken from, with the fifth time point representing the time-series run (t_grouped_). The x-axis shows the four sample sizes taken from each source population. Each column represents the five tested admixture times (t_admix_), using a gene-flow rate of 10%, and each row the results for the four methods. Each square shows the average across all 10 samples from the admixed population and across all 10 replicates. Higher sensitivity estimates represent higher true positive rates.

**Fig. S21:**
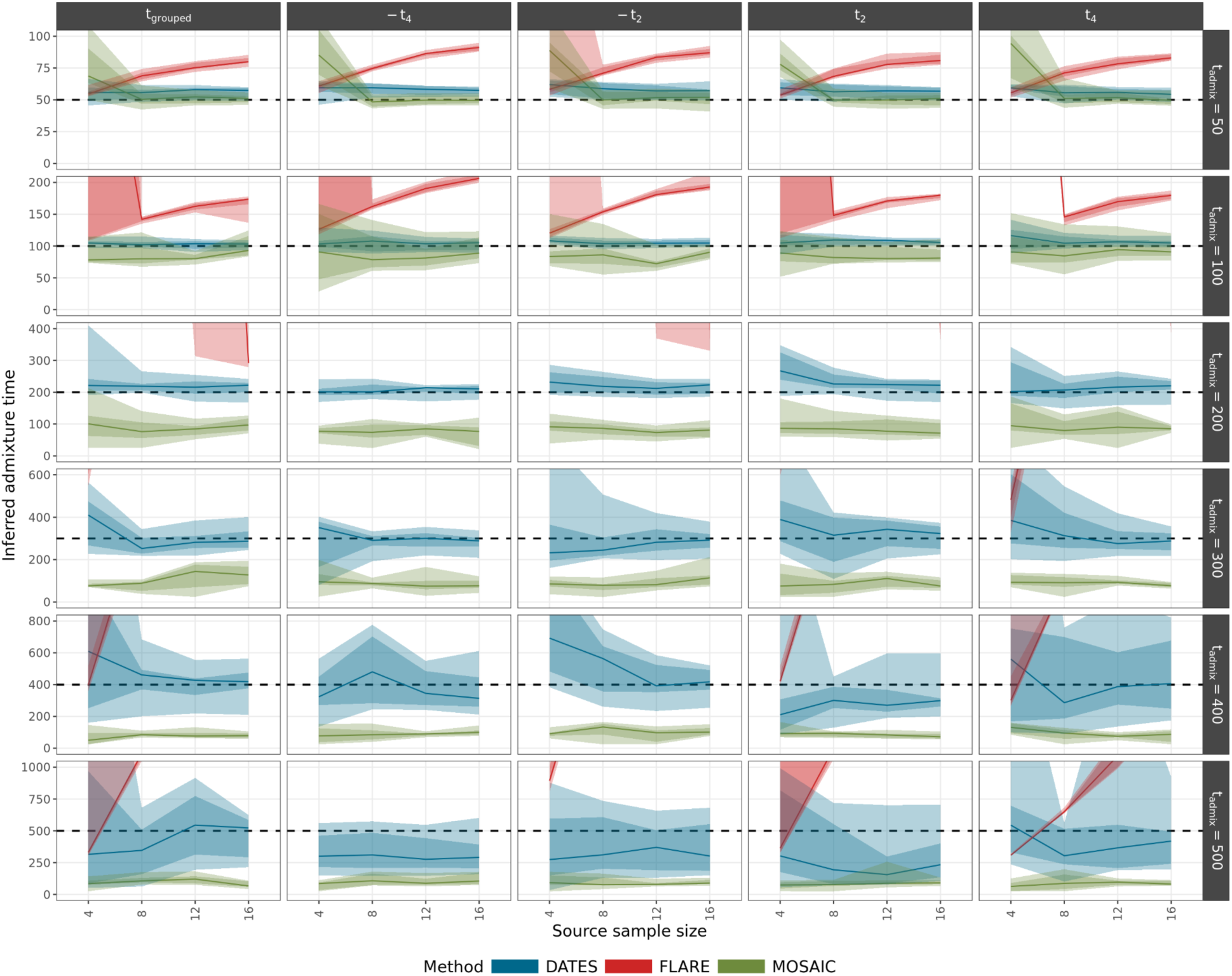
Inferred admixture times (y axis) under a subset of demographic parameters from DATES (blue), FLARE (red) and MOSAIC (green), using a merged time series dataset of sources sampled from four time points (t_sampling_ : t_-4_=t_admix_×0.2, t_-2_=t_admix_×0.6, t_2_=t_admix_×1.4, t_4_=t_admix_×1.8). The x-axis shows the four sample sizes taken from each of the two ancestries. Each column the four time points at which the source samples were taken from, with the first time point representing the time-series run (t_grouped_), and each row one of the six tested admixture times (t_admix_). All results are shown for a gene-flow rate of 30%. The dotted black line indicates the true admixture time. The shaded regions in each ribbon show the 50% and 80% quantiles and the line represents the median. Estimates which fall outside of the range [0, 2×t_admix_] are not shown.

**Fig. S22:**
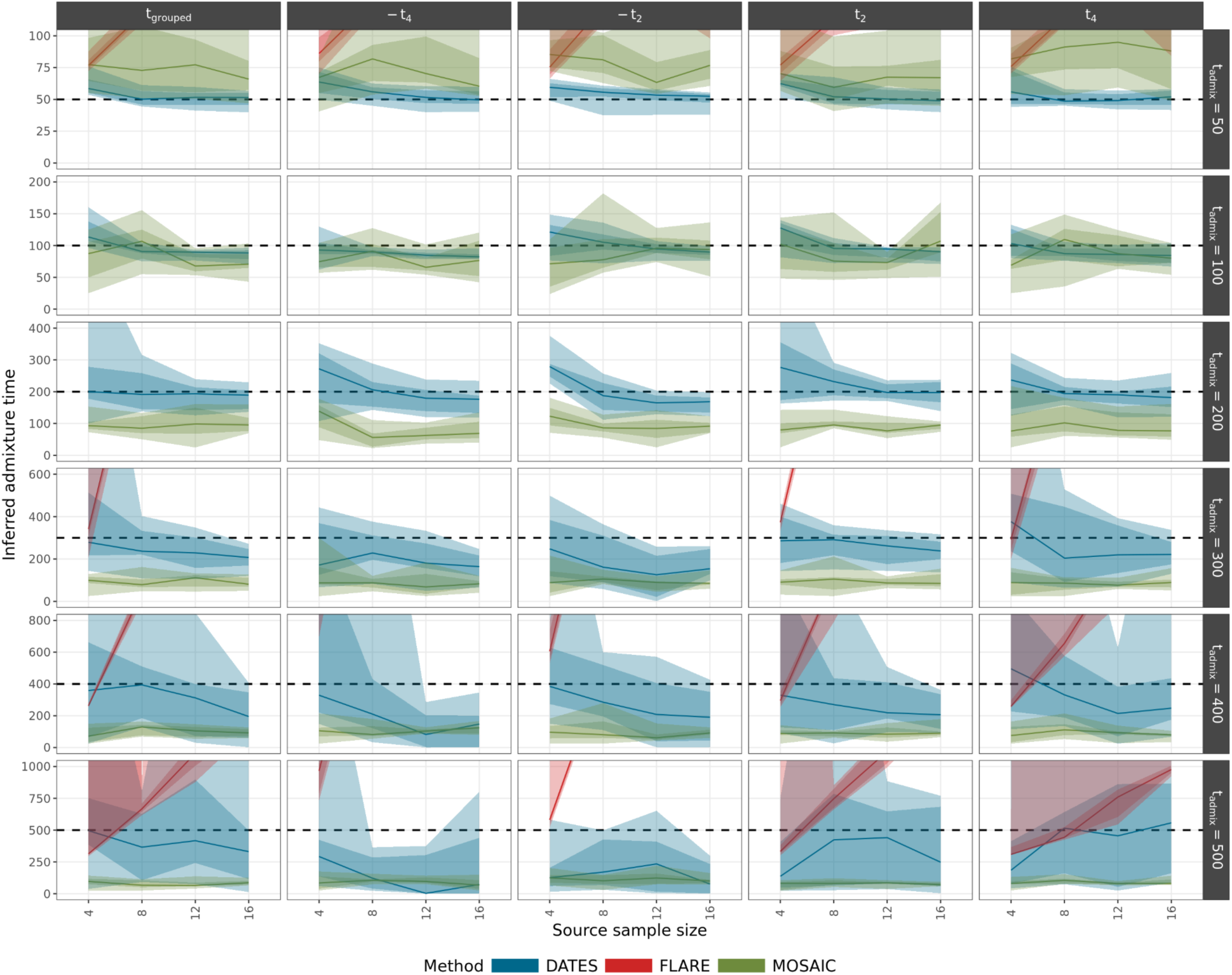
Inferred admixture times (y axis) under a subset of demographic parameters from DATES (blue), FLARE (red) and MOSAIC (green), using a merged time series dataset of sources sampled from four time points (t_sampling_ : t_-4_=t_admix_×0.2, t_-2_=t_admix_×0.6, t_2_=t_admix_×1.4, t_4_=t_admix_×1.8). The x-axis shows the four sample sizes taken from each of the two ancestries. Each column the four time points at which the source samples were taken from, with the first time point representing the time-series run (t_grouped_), and each row one of the six tested admixture times (t_admix_). All results are shown for a gene-flow rate of 10%. The dotted black line indicates the true admixture time. The shaded regions in each ribbon show the 50% and 80% quantiles and the line represents the median. Estimates which fall outside of the range [0, 2×t_admix_] are not shown.

